# Immune Signatures of SARS-CoV-2 Infection Resolution in Human Lung Tissues

**DOI:** 10.1101/2024.03.08.583965

**Authors:** Devin Kenney, Aoife K. O’Connell, Anna E. Tseng, Jacquelyn Turcinovic, Maegan L. Sheehan, Adam D. Nitido, Paige Montanaro, Hans P. Gertje, Maria Ericsson, John H. Connor, Vladimir Vrbanac, Nicholas A. Crossland, Christelle Harly, Alejandro B. Balazs, Florian Douam

## Abstract

While human autopsy samples have provided insights into pulmonary immune mechanisms associated with severe viral respiratory diseases, the mechanisms that contribute to a clinically favorable resolution of viral respiratory infections remain unclear due to the lack of proper experimental systems. Using mice co-engrafted with a genetically matched human immune system and fetal lung xenograft (fLX), we mapped the immunological events defining successful resolution of SARS-CoV-2 infection in human lung tissues. Viral infection is rapidly cleared from fLX following a peak of viral replication, histopathological manifestations of lung disease and loss of AT2 program, as reported in human COVID-19 patients. Infection resolution is associated with the activation of a limited number of hematopoietic subsets, including inflammatory monocytes and non-canonical double-negative T-cells with cytotoxic functions, which are highly enriched in viral RNA and dissipate upon infection resolution. Activation of specific human fibroblast and endothelial subsets also elicit robust antiviral and monocyte chemotaxis signatures, respectively. Notably, systemic depletion of human CD4+ cells, but not CD3+ cells, abrogates infection resolution in fLX and induces persistent infection, supporting evidence that peripheral CD4+ monocytes are important contributors to SARS-CoV-2 infection resolution in lung tissues. Collectively, our findings unravel a comprehensive picture of the immunological events defining effective resolution of SARS-CoV-2 infection in human lung tissues, revealing markedly divergent immunological trajectories between resolving and fatal COVID-19 cases.

## INTRODUCTION

Coronavirus disease 2019 (COVID-19) is a respiratory disease that has swept the world since its emergence in the Wuhan province of China in late 2019. The etiologic agent of COVID-19, the Severe Acute Respiratory Syndrome Coronavirus 2 (SARS-CoV-2) is a plus-sense, enveloped RNA virus that targets the epithelium of the respiratory tract. Infection results in varying severities of COVID-19, with most cases being mild to asymptomatic. The onset of severe disease is associated with aberrant immune responses (e.g., excessive pulmonary infiltration of myeloid cells, inflammasome-activated monocytic cells, macrophage exacerbated inflammation) and severe lung injury (e.g., lung consolidation, diffuse alveolar damage (DAD), and thrombosis) (*1-9*).

A large number of human studies have been instrumental in unraveling cellular and molecular processes driving severe COVID-19 disease in infected tissues, particularly in the respiratory tract, using autopsy samples (*10-13*). In parallel, human studies of resolving COVID-19, including controlled human challenge studies, have leveraged peripheral blood and nasopharyngeal samples to identify human signatures of effective infection resolution and mild disease (*14-16*). Notably, this includes individuals with specific HLA haplotypes (*15*), evidence of previous coronavirus exposure (*15, 17, 18*), rapid nasopharyngeal immune infiltration (*14*), and non-productive infection of nasopharyngeal T-cells and macrophages (*14*). However, tissue-specific human immunological processes associated with protection, such as extravasation of recruited immune lineages and their differentiation processes, have remained elusive due to the ethical considerations associated with tissue sampling of individuals with mild disease and human challenge models. Although large and small animal models of COVID-19 are available, the high-cost and limited reagent availability associated with non-human primate models, and the large divergence between rodent and human immune systems (*19, 20*) further underscore the need for additional models capable of recapitulating human protective immune responses to SARS-CoV-2 infection.

Mice engrafted with human fetal lung xenograft (fLX) support infection by multiple human respiratory viruses, including human cytomegalovirus (HCMV), Middle East respiratory syndrome coronavirus (MERS-CoV) and SARS-CoV-2 (*21-23*). Upon co-engraftment with a human immune system (HIS), these animals also mount lung-resident human immune responses against these pathogens (*21, 23-25*). Recently, our group reported that mice engrafted with fLX are highly susceptible to SARS-CoV-2 infection and lung tissue damage and support persistent viral infection (*23*). However, co-engraftment of fLX and HIS in a xenorecipient strain supporting enhanced myelopoiesis (i.e., HNFL mouse model) rapidly blunted viral infection and prevented widespread acute viral replication across fLX, resulting in protection from histopathology of fLX (*23*). Our findings unraveled a human macrophage antiviral signature as a correlate of such rapid protection against SARS-CoV-2 infection. However, how the human immune system mobilizes and resolves infection upon extensive viral replication within human lung tissues, a context that more likely describes mild cases of SARS-CoV-2 infection, has remained elusive.

In this study, we leverage previously described immunodeficient mice engrafted with a human fetal lung xenograft (fLX) and a genetically matched human immune system, fetal liver and thymus (BLT-L mice) (*21*) to conduct the first comprehensive mapping of human immunological correlates of resolution of acute SARS-CoV-2 infection in human lung tissues. Consistent with previous reports (*24, 25*), BLT-L mice are permissive to SARS-CoV-2 infection following direct viral inoculation into the fLX. Infection swiftly resolves by 6 days post-infection (dpi) following an early viral replication peak at 2 dpi. Acute viral infection is associated with lung histopathological damage and loss of AT2 program, as described in COVID-19 patients (*1*). Notably, acute infection is also defined by the emergence of a limited set of hyper-activated hematopoietic subsets, including inflammatory monocytes and T-cells expressing genes canonically found in macrophages. Of these, inflammatory monocytes appear to mount the most robust antiviral responses and are highly enriched in viral RNA before phasing out from the tissues after resolution. No inflammasome activation in these cells was observed despite enrichment in viral RNA. This contrasts with reports from fatal COVID-19 cases (*26*) and evidence suggesting that monocyte and macrophage infections in vitro and in vivo are associated with inflammasome activation (*27, 28*).

Specific fibroblast and endothelial cell subsets also contribute to antiviral responses and hematopoietic chemotaxis, respectively, underscoring how crosstalk between hematopoietic and non-hematopoietic lineages likely contributes to infection resolution. At 12 dpi, the immune landscape in fLX is characterized by an increase in CD4+ patrolling monocytes, conventional dendritic cells and CD206+ interstitial macrophages (IM). Notably, systemic depletion of human CD4+ cells, but not human CD3+ cells, abrogates SARS-CoV-2 clearance in fLX and causes persistent infection. These findings underscore CD4+ circulating monocyte infiltration as a likely feature that defines SARS-CoV-2 infection resolution.

Collectively, our work sheds light on a unique set of immunological events associated with SARS-CoV-2 resolution in human lung tissues which dramatically contrasts with the immune trajectories reported in fatal COVID-19 cases. This work opens avenues for mechanistically dissecting the respective contributions of the hematopoietic and non-hematopoietic subsets identified in this study, which could inform the development of immunotherapies against viral respiratory diseases.

## RESULTS

### BLT-L mice effectively clear SARS-CoV-2 infection following acute viral replication

Previous work from our laboratory (*23*) and others (*21, 24, 25*) have shown that immunodeficient mice can successfully be engrafted with human fetal lung xenograft (fLX) alone or in combination with a human immune system (HIS). In this study, we leveraged a previously reported mouse model co-engrafted with fetal liver and thymus as well as with human hematopoietic stem cells (HSC) and fLX (BLT-L mice) (*21*). Fetal liver and thymus were engrafted under the renal capsule of adult NOD.Cg-*Prkdc^scid^ Il2rg^tm1Wjl^*/SzJ (NSG) mice (12-16 weeks old) prior to intravenous HSC injection. A piece of fetal lung tissue was subcutaneously engrafted on the flank of each animal, as described previously (*21*) **(Fig. 1A)**. To determine the susceptibility of BLT-L mice to SARS-CoV-2 infection and their ability to effectively clear infection, BLT-L mice were inoculated with SARS-CoV-2 (2019-nCoV/USA_WA1/2020) via intra-graft injection. We used a viral dose (10^6^ PFU) that we previously established to drive robust and persistent infection in fLX of immunodeficient mice not engrafted with a HIS (*23*) (**Fig. 1B)**.

**Figure 1.**
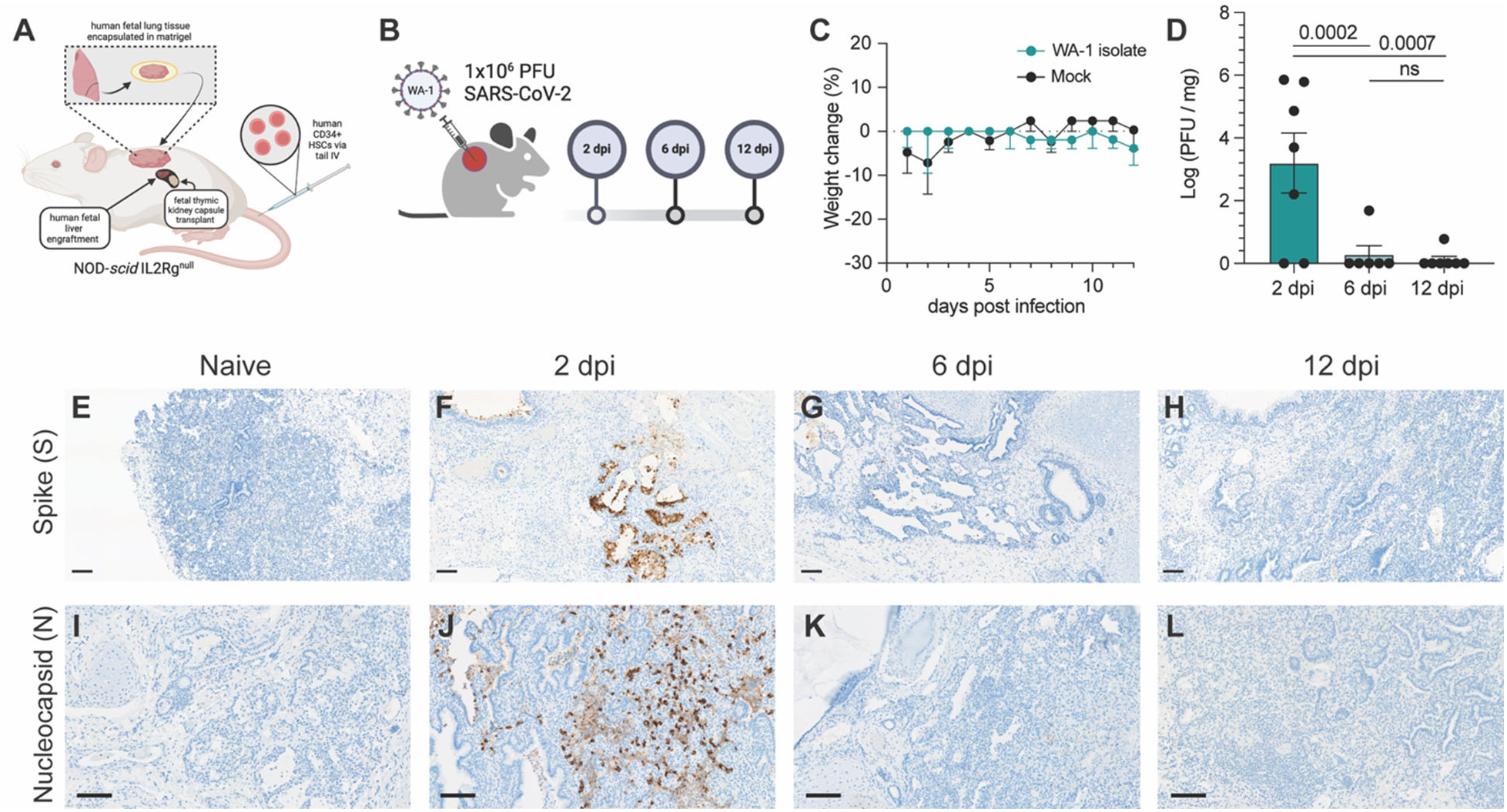
BLT-L mice effectively resolve SARS-CoV-2 infection. **(A)** Representative schematic of BLT-L mice. **(B)** Fetal lung xenografts (fLX) of BLT-L mice were infected with 1×10^6^ PFU of SARS-CoV-2 WA-1 isolate via subcutaneous intra-graft injection. **(C)** BLT-L mice were monitored for weight change over the course of 12 days post-infection (dpi). **(D)** Viral titer (log(PFU/mg)) in fLX at 2, 6, and 12 dpi as determined by plaque assay. **(E-H)** Immunohistochemistry for SARS-CoV-2 S and **(I-L)** N protein were performed on naïve fLX **(E,I)** and fLX at 2 dpi **(F,J)**, 6dpi **(G,K),** and 12 dpi **(H,L)**. Scale bar = 100 μM. Data are representative of two or three independent experiments. n = 6 -10 per timepoint. Error bars represent mean *±* standard error of the mean. *One-way ANOVA analysis was performed. p-values are indicated. n.s. = non-significant*.

Throughout the course of infection, mice did not display any weight loss (**Fig. 1C)** or clinical signs of disease such as lethargy or lack of responsiveness (data not shown). To assess lung histopathology and viral titers longitudinally, fLX were collected at 2, 6, and 12 dpi. Plaque assay was performed on fLX homogenates to determine viral titers. A significant amount of infectious viral particles could be recovered from fLX at 2 dpi (3.20 ± 2.52 log([PFU/mg of tissue]), but not at 6dpi (0.281 ± 0.688 log[PFU/mg of tissue]) and 12 dpi (0.111 ± 0.293 log[PFU/mg of tissue]) (**Fig. 1D**). These data demonstrated a peak of viral infection at 2 dpi, prior to resolution of infection by 6 dpi. Immunohistochemistry (IHC) for SARS-CoV-2 Spike (S) protein revealed infection was mainly found in the alveolar epithelium of fLX at 2 dpi (**Fig. 1E,F**). Consistent with viral quantification, most viral antigen was cleared by 6 dpi and became undetectable by 12 dpi (**Fig.1G,H**). SARS-CoV-2 S was primarily detected in the alveolar and bronchiole epithelium along with necrotic cellular debris, consistent with the primary cell targets of SARS-CoV-2 (**Fig.1E**). These findings were confirmed through IHC for SARS-CoV-2 nucleoprotein (N) at 2, 6 and 12 dpi (**Fig. 1I-L**) and transmission electron microscopy (TEM) imaging of fLX at 2 dpi **(Fig. S1A-C)**. Notably, TEM substantiated evidence of productive fLX infection, as indicated by the presence of viral particles in the cytosol of epithelial cells and budding/invagination of virions (**Fig. S1A-C**).

Next, we examined histopathological phenotypes associated with active and resolved infection. Interpretation of hematoxylin and eosin (H&E) staining illustrated denuding of pneumocytes, neutrophil infiltration, edema, hemorrhage, thrombosis, and pneumocyte necrosis, which correlated with sites of infection at 2 dpi (**Fig. 2A-E**). No major signs of histopathological lung damage were observed at 6 and 12 dpi compared to naïve fLX, indicating that fLX can mount repair mechanisms upon resolution of infection (**Fig. 2F-G**). A previously described semi-quantitative histopathological scoring system (*23*) provided statistical confirmation for a significant increase in lung pathology at 2 dpi, which was no longer apparent at 6 (**Fig. 1H**) or 12 dpi. Of note, minor lung pathology was observed at baseline (naïve), likely reflecting limited graft vs. host disease. Together, infection of fLX of BLT-L mice recapitulates many important hallmarks of acute SARS-CoV-2 infection, including viral replication and histopathological manifestations of disease, prior to effective viral clearance and lung tissue repair.

**Figure 2.**
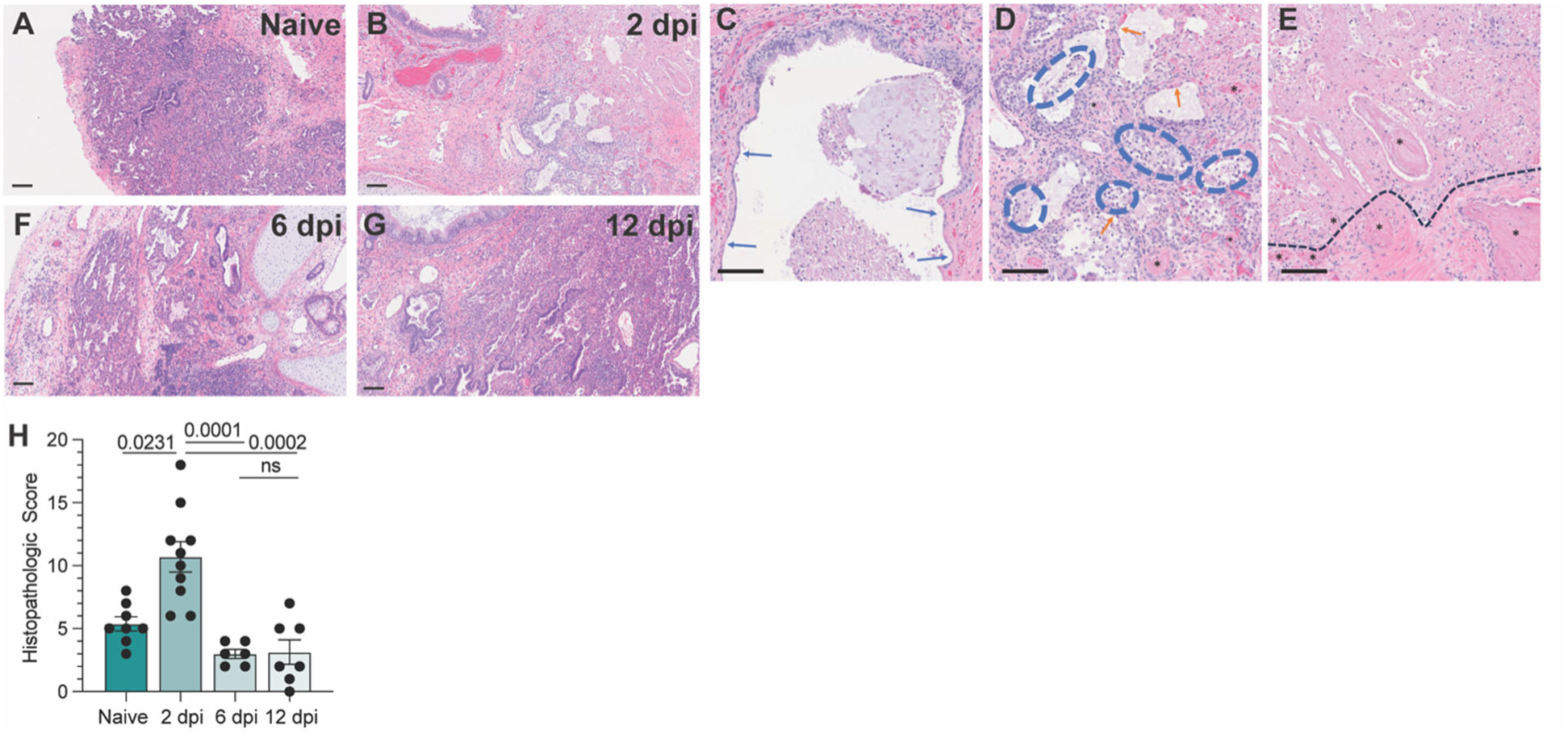
BLT-L mice resolve histopathological damage in fLX. **(A-E)** Hematoxylin and eosin staining was performed on naïve fLX **(A)** and fLX at 2 dpi **(B-E)**, 6dpi **(F),** and 12 dpi **(G)**. Scale bar = 100 μM. **(C)** Bronchiolar attenuation and denuding (blue arrows) correlates directly with the presence of SARS-CoV-2 Spike protein. **(D)** Alveolar spaces are multifocally filled with necrotic debris admixed with neutrophils and edema (blue hashes). Type II pneumocytes (ATII) are often denuded or attenuated in areas of inflammation that correlates directly with the presence of SARS-CoV-2 Spike protein. Fibrin thrombi are routinely observed in neighboring parenchyma (black asterisks). **(E)** Coagulative necrosis as evidenced by loss of cellular detail and generalized eosinophilia (above the black hashed line) with numerous regional fibrin thrombi (black asterisks). Although SARS-CoV-2 Spike protein is not observed in the area of coagulative necrosis viral antigen is located within adjacent tissue. **(H)** Histopathological scoring of naïve fLX and fLX at 2, 6, and 12 dpi. Data are representative of two or three independent experiments. n = 6 -10 per timepoint. Error bars represent mean *±* standard error of the mean. *One-way ANOVA analysis was performed. p-values are indicated. n.s. = non-significant*.

### Humoral responses do not drive SARS-CoV-2 clearance in BLT-L mice despite evidence of Spike selective pressure

We first asked whether SARS-CoV-2 infection resolution was driven by human neutralizing humoral responses. Consistent with the rapid clearance of infectious viral particles by 6 dpi and the known caveat that humanized mice mount limited humoral responses (29), there were no detectable neutralizing antibodies in serum collected at 2 or 12 dpi (**Fig. S2A, B**). However, interestingly, genomic sequencing of virus isolated from fLX at 2 dpi revealed the selection of two stable mutations in 75% of fLX (**Fig. S2C**, **Table S1**). Both mutations were located in the Spike N-terminal domain (NTD): an insertion (216KLRS) and a non-synonymous mutation (R245H), neither of which were present in the inoculum (**Fig. S2C**, **Table S1**). Interestingly, these two mutations were found together in 100% of the viral sequences, suggesting potential co-evolution (**Fig. S2D**). They have also been reported as positively selected in the context of sub-optimal neutralizing antibody concentration (216KLRS) (*29*) or the context of cross-species adaptation (216KLRS and R245H) (*30*). Despite lacking humoral responses, these findings reveal that BLT-L mice can recapitulate host-pathogen interactions that drive SARS-CoV-2 evolution and host adaptation.

### SARS-CoV-2 infection remodels the human lung cellular environment

To comprehensively map the fLX responses associated with resolution of SARS-CoV-2 infection, we performed single-cell RNA sequencing (scRNA-seq) on fLX from naïve BLT-L mice and at 2 and 12 dpi. While most cells detected in fLX by scRNA-seq were human, a minor population of mouse cells was detected, which were excluded from downstream analysis (**Fig. S3A-C**). Initial analysis of the human compartment revealed diverse hematopoietic (T-cells/innate lymphoid cells (ILC), B cells, myeloid cells and mast cells) and non-hematopoietic lineages (ciliated and non-ciliated epithelial, endothelial, mesenchymal cells and chondrocytes) in both naïve and infected fLX (**Fig. 3A-C and Fig. S3D,E**).

**Figure 3.**
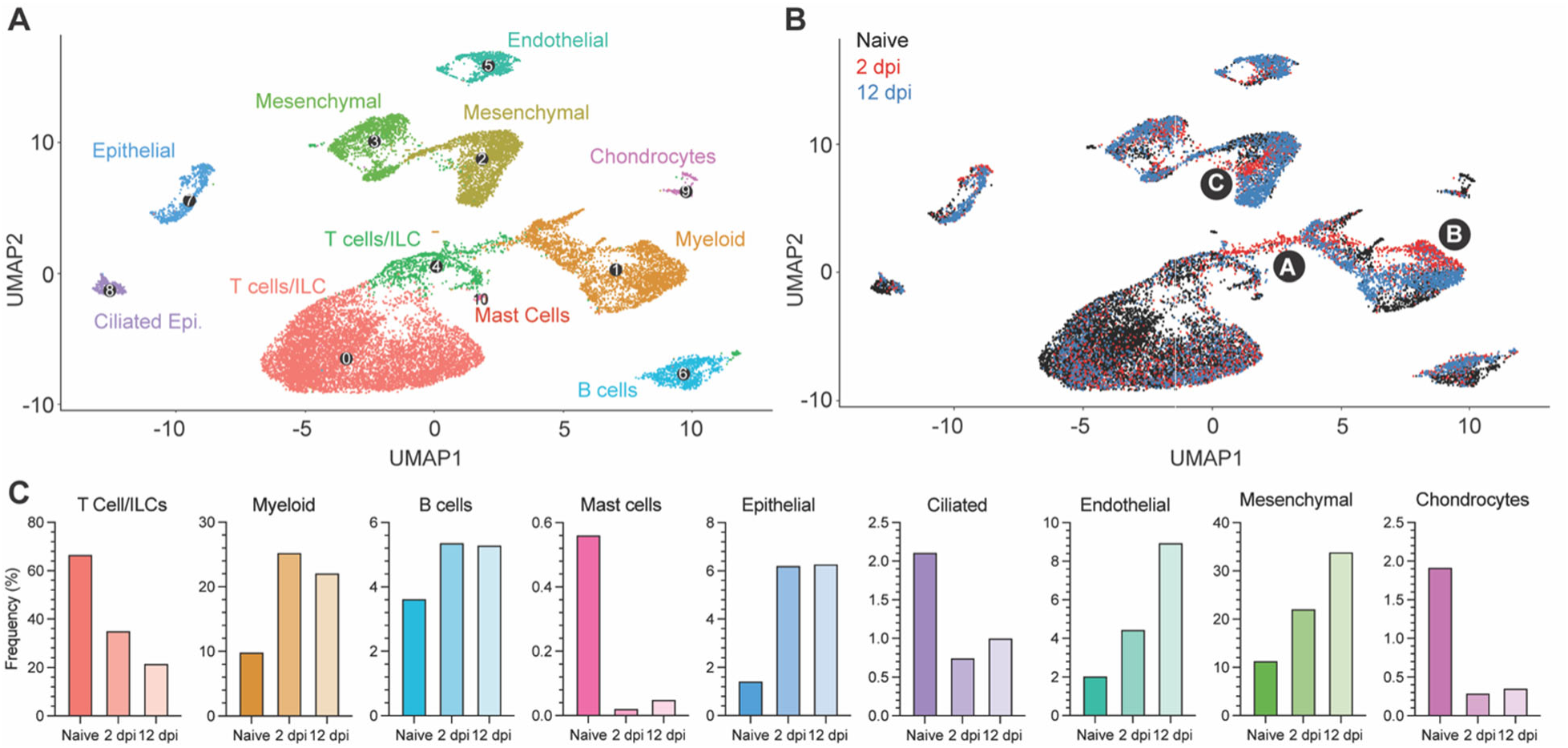
Cellular remodeling and viral dynamics in fLX upon SARS-CoV-2 infection. Single cell RNA sequencing analysis was performed on single cell suspensions of naïve fLX and fLX at 2 and 12 dpi. Naïve n=3 fLX (9,605 cells), 2 dpi n=2 fLX (4,405 cells), and 12 dpi n = 2 fLX (5,857 cells). **(A)** UMAP plot clustering of the human cell compartment of naïve fLX and fLX at 2 and 12 dpi. **(B)** Temporal annotation of the human clusters on the UMAP plot: naïve (black), 2 dpi (red), 12 dpi (blue). Sub-clusters unique to 2 dpi are annotated by A (T cell compartment), B (myeloid compartment) and C (mesenchymal compartment). **(C)** Frequency of each cell compartment determined by single cell RNA-sequencing.

T-cells and innate lymphoid cells (ILC) have the same transcriptional programs, express TCR genes, and play similar effector functions (*31*). Their shared features render them very challenging to distinguish by scRNAseq in small datasets and without TCR sequencing information (*32*). We thus considered them together as part of a T-cell/ILC population. T-cell/ILC frequency dramatically decreased in fLX upon infection (naïve, 66.8%; 2 dpi, 35.3%; 12 dpi, 21.8%), which is consistent with evidence that COVID-19 can induce lymphoid depletion(*33, 34*) (**Fig. 3B,C**). Notably, lymphopenia was not observed in the peripheral blood of BLT-L mice, suggesting lymphocyte depletion in fLX was not directly attributed to declining circulating peripheral lymphocytes (**Fig. S3F)**. In contrast, myeloid (naïve, 9.94%; 2 dpi, 25.3%; 12 dpi, 22.2%) and B-cell subsets (naïve, 3.64%; 2 dpi, 5.38%; 12 dpi, 5.31%) relatively expanded upon infection (**Fig. 3B,C**). The epithelial, endothelial and mesenchymal cell frequencies increased upon infection, while mast cell, ciliated cell and chondrocyte frequencies decreased (**Fig. 3B,C**). Notably, most human clusters showed temporal segregation between naïve fLX and fLX at 2 and 12 dpi (**Fig. 3B and Fig. S3E**; naïve: black subclusters; 2 dpi: red subclusters and 12 dpi: blue subclusters). This suggests that infection alters the transcriptional state of many cell types and/or drives the emergence of novel cell subsets. Notably, acute infection at 2 dpi led to the emergence of distinct T, myeloid, and mesenchymal cell populations, labeled as subclusters A, B, and C, respectively (**Fig. 3B**). Resolved infection (12 dpi) was associated with the emergence of transcriptomically distinct epithelial, endothelial, mesenchymal, B-cell and myeloid subclusters; while we did not observe the emergence of transcriptomically divergent subclusters within the T-cell lineages at that time point. Temporal separation of several human lineages suggests a two-step tissue remodeling process during infection involving i) an initial antiviral phase mediated by a limited set of human subpopulations and ii) a tissue repair phase involving a broader range of human subpopulations.

Next, we wished to utilize scRNAseq to determine the cellular compartments enriched in viral RNA. We found that viral RNA was mainly within three major lineages: mesenchymal, T-cell/ILCs and myeloid (**Fig.4A,B**). AT2 and the overall epithelial compartment were not significantly enriched in viral RNA despite histological evidence of epithelial infection in fLX at 2 dpi (**Fig. 4A and 1F,J**). Previous evidence that SARS-CoV-2 induces significant cytopathic damage in AT2 cells (*35*) may suggest that infected AT2 cells are lost when undergoing the scRNAseq pipeline. Within the mesenchymal cluster, viral reads were limited and distributed sporadically without significant enrichment within a given subcluster (**Fig. 4B,C**). However, viral RNA was strongly enriched within the 2 dpi-specific populations in the myeloid and T-cell clusters, previously labeled as clusters A and B (**Fig. 4A,B**), warranting further investigations.

**Figure 4.**
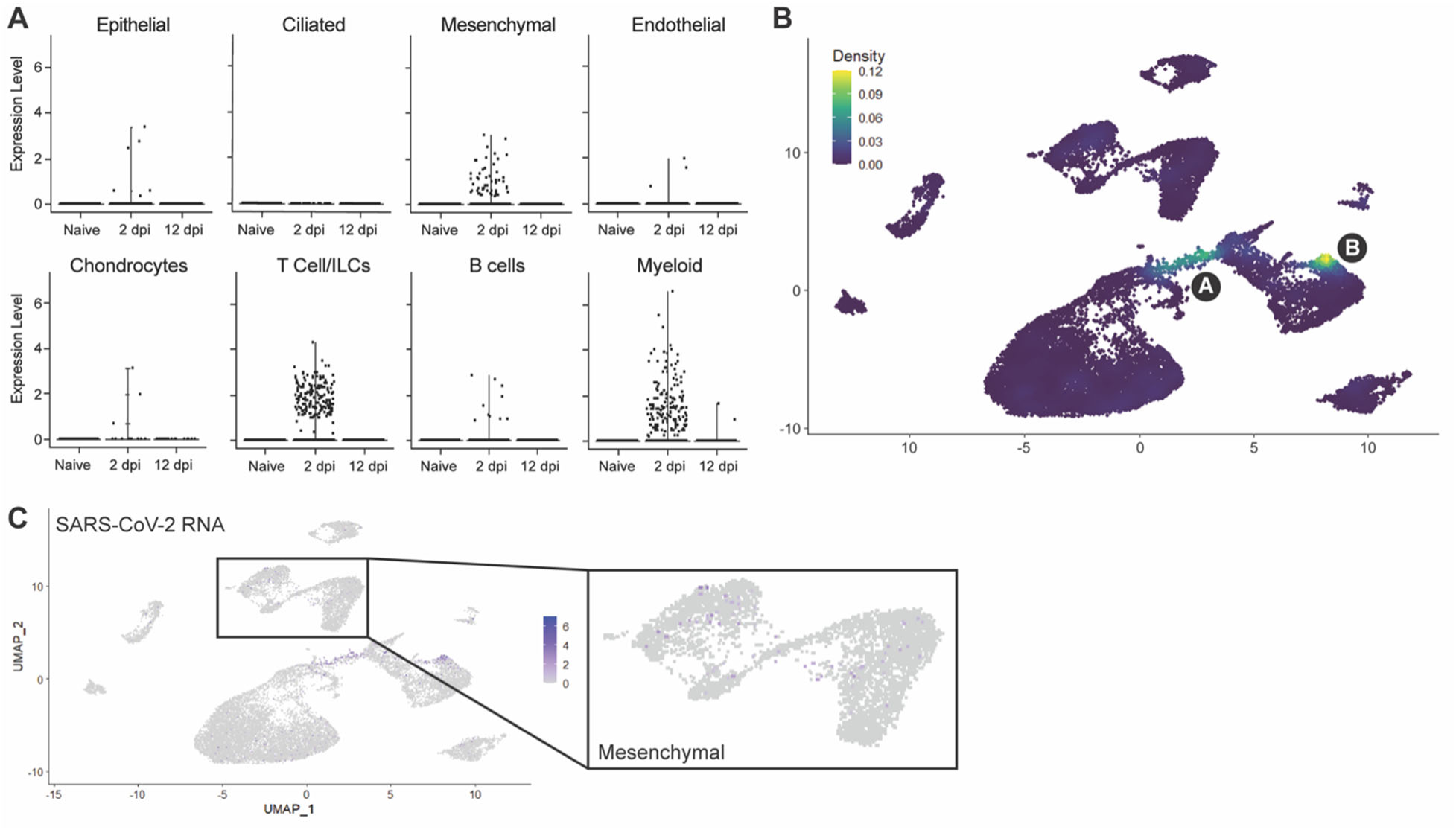
Cellular compartmentalization of viral RNA during infection. **(A)** Dot plots displaying the expression of SARS-CoV-2 viral RNA transcripts identified in scRNA-seq. **(B)** UMAP plot showing the distribution of SARS-CoV-2 viral RNA transcripts by density across all human cell clusters and time points. **(C)** UMAP plot showing the distribution of SARS-CoV-2 viral RNA reads across all human cell clusters and time points. Inset showing distribution of viral RNA reads in the mesenchymal cluster.

### Human lung epithelium signatures upon SARS-CoV-2 infection recapitulate features of COVID-19

Subclustering of the human epithelial compartment revealed subclusters of airway basal and secretory cells, alveolar type 1 (AT1) and type 2 (AT2) cells, and serous cells (**Fig. 5A**). The dynamic changes of the epithelial compartment upon infection were consistent with COVID19 human studies (*2, 5, 36*). We noted a relative reduction of the AT2 compartment at 2 dpi (25.6%) compared to naïve mice (73.1%). The AT2 compartment was partially restored at 12 dpi (49.1%) (**Fig. 5B**). The frequency of basal airway cells followed a reversed trend, indirectly reflecting the disruption of the AT2 compartment. The size of the AT1 compartment showed a relative increase at 2 dpi (9.89% compared to 8.00% in naïve fLX), consistently with the ability of AT2 cells to differentiate into AT1 upon lung damage (*37*) before returning to minimal relative levels as the AT2 compartment re-expanded following infection resolution.

**Figure 5.**
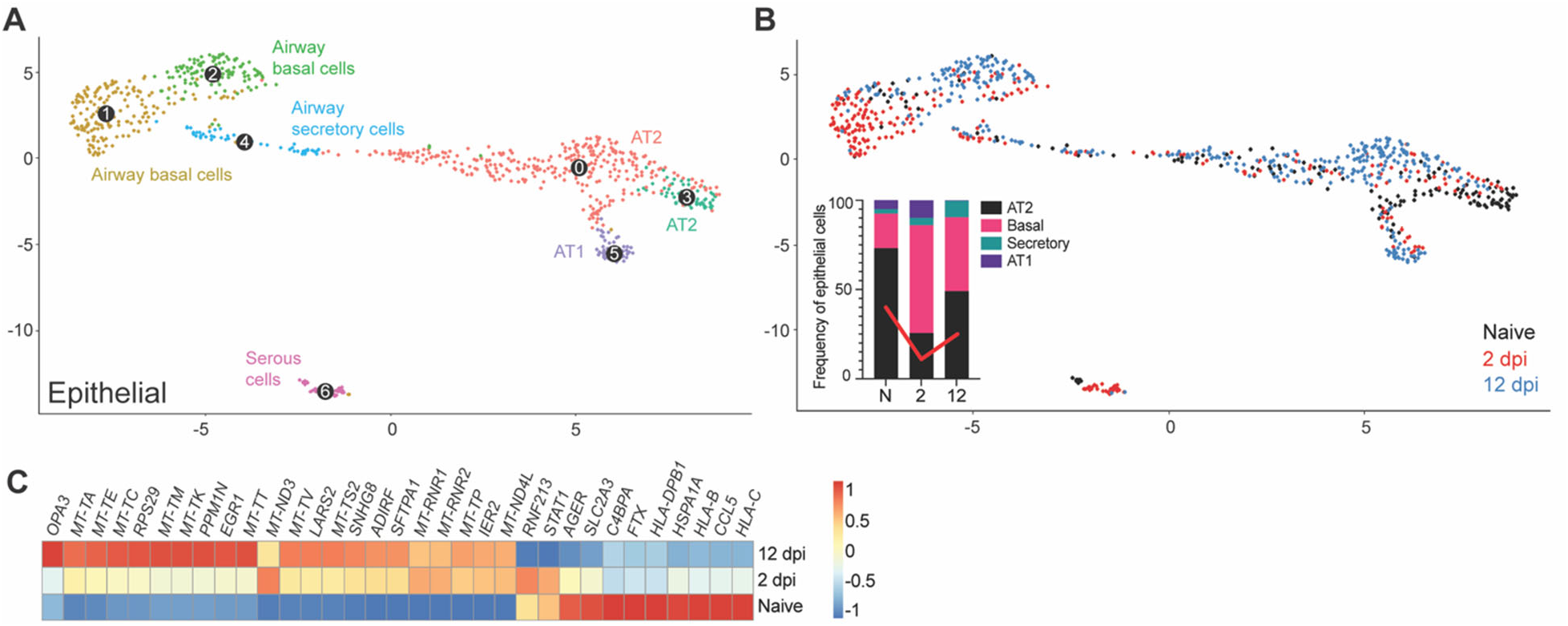
Remodeling of the epithelial compartment of fLX during infection. **(A)** UMAP plot sub-clustering of the human epithelial compartment in fLX across all time points. **(B)** Temporal annotation of the human epithelial sub-clusters on the UMAP plot: naïve (black), 2 dpi (red), 12 dpi (blue). The inset graph indicates the change in frequency of each compartment (AT2; black, Basal; pink, Secretory; teal, and AT1; purple) within the epithelial compartment. Red line indicates the change in the AT2 compartment over time. **(C)** Heatmap displaying the relative expression (from -1 to 1) of the top differentially regulated genes over the course of infection within the AT2 compartment.

The AT2 compartment displayed the most robust remodeling upon viral infection across the entire epithelial compartment, with 2 dpi and 12 dpi AT2 subclusters showing distinctive transcriptomic signatures compared to naïve AT2 cells (**Fig. 5C**). Interestingly, AT2 cells did not exhibit robust antiviral responses upon acute infection (**Fig. 5C**). However, they displayed gene signatures previously reported during and after SARS-CoV-2 infection including, downregulation of MHC genes (*HLA-C, HLA-B, and HLA-DPB1)* and *CCL5* (a cytokine associated with immune recruitment during respiratory infection) (*6*), upregulation of immunomodulatory genes regulating inflammation and cell stress (*SFTPA1*, *EGR1*), and elevated expression of genes associated with mitochondrial dysfunctions and increased oxidative stress responses (i.e., *LARS2*). (**Fig. 5C**). Activation of tissue repair mechanism (*SNHG8*) was also observed, further underscoring evidence of physiologically relevant AT2 response to viral infection (**Fig. 5C**). The absence of detectable viral RNA in AT2 cells was in line with the lack of antiviral responses. Such a lack of infected AT2 cells in our scRNAseq data is likely the reflection of their vulnerability and death during infection, given histological evidence of epithelial infection (**Fig. 1F, J**) and the marked reduction of the AT2 compartment in fLX upon infection (**Fig. 5B**).

Collectively, the epithelial compartment of fLX of BLT-L mice recapitulates several previously reported features of COVID-19 in humans, including loss of AT2 program, downregulation of immune genes and antigen presentation, and activation of tissue repair mechanisms.

### SARS-CoV-2 infection resolution is associated with the emergence of a viral RNA-enriched double-negative T-cell subcluster with cytotoxic functions

We first aimed to characterize the T-cell signatures to SARS-CoV-2 infection resolution, and with that, the features of our viral RNA-enriched, 2 dpi-specific, T-cell/ILC subcluster A. T-cell/ILC responses to infection were mainly limited to this specific viral RNA-associated subcluster (**Fig. 6A-D**), which segregated very distinctively from other CD3+ T-cell lineages, including canonical CD4+ and CD8+ T-cells (Subcluster 6; **Fig. 6A-C**). Using doublet discriminators and assessing viral RNA abundance; we determined that this subcluster was not a doublet and displayed low levels of *CD4* and *CD8* transcripts, suggesting a double-negative profile, and elevated levels of mitochondrial transcripts **(Fig. S4A,B**). Most notably, it was uniquely defined by the expression of several transcripts known to be enriched in macrophages (*MARCO*, *TIMP1*, *LYZ*), fibroblasts (*MGP*, *CALD1*, *COL1A*), or both (*A2M*, *IFITM3*) but not in T-cells (**Fig. 6E and Fig. S4C**). This subcluster also exhibited evidence of cytotoxic function through the expression of *GZMA*, *GZMB*, and *GNLY* transcripts, albeit expression was also detected in CD8+ T-cells and activated tissue-resident memory T-cells (TRM) (Subclusters 1,3,4, **Fig. 6E**). Notably, a subpopulation of double-negative T-cells sharing some of these cytotoxic markers has been previously reported in mouse spleen(*38*). Given the proximity of this subcluster with myeloid lineages (**Fig. 3A,B**) and evidence of expression of macrophage-enriched transcripts, we referred to this subcluster as myeloid-like double-negative T-cells (mDNT cells). mDNT cells also exhibited upregulation of key gene pathways related to COVID-19, SARS-CoV-2 cell-intrinsic immune responses, cellular cytotoxicity (e.g., degranulation) and cell death, consistently with a role for this subset to serve as a robust primary responder to viral infection (**Fig. 6D**). Of note, a recent human challenge study reported that self-resolving SARS-CoV-2 infection is associated with non-productive infection of human nasopharyngeal T-cells(*14*). In contrast, in lung autopsy samples of fatal COVID-19 cases, viral RNA was enriched in myeloid cells but was not detected in T-cells(*39*). Collectively, our findings underscore that the emergence of viral RNA-enriched T cell populations displaying myeloid-like features in infected lung tissues is associated with effective SARS-CoV-2 infection resolution.

**Figure 6.**
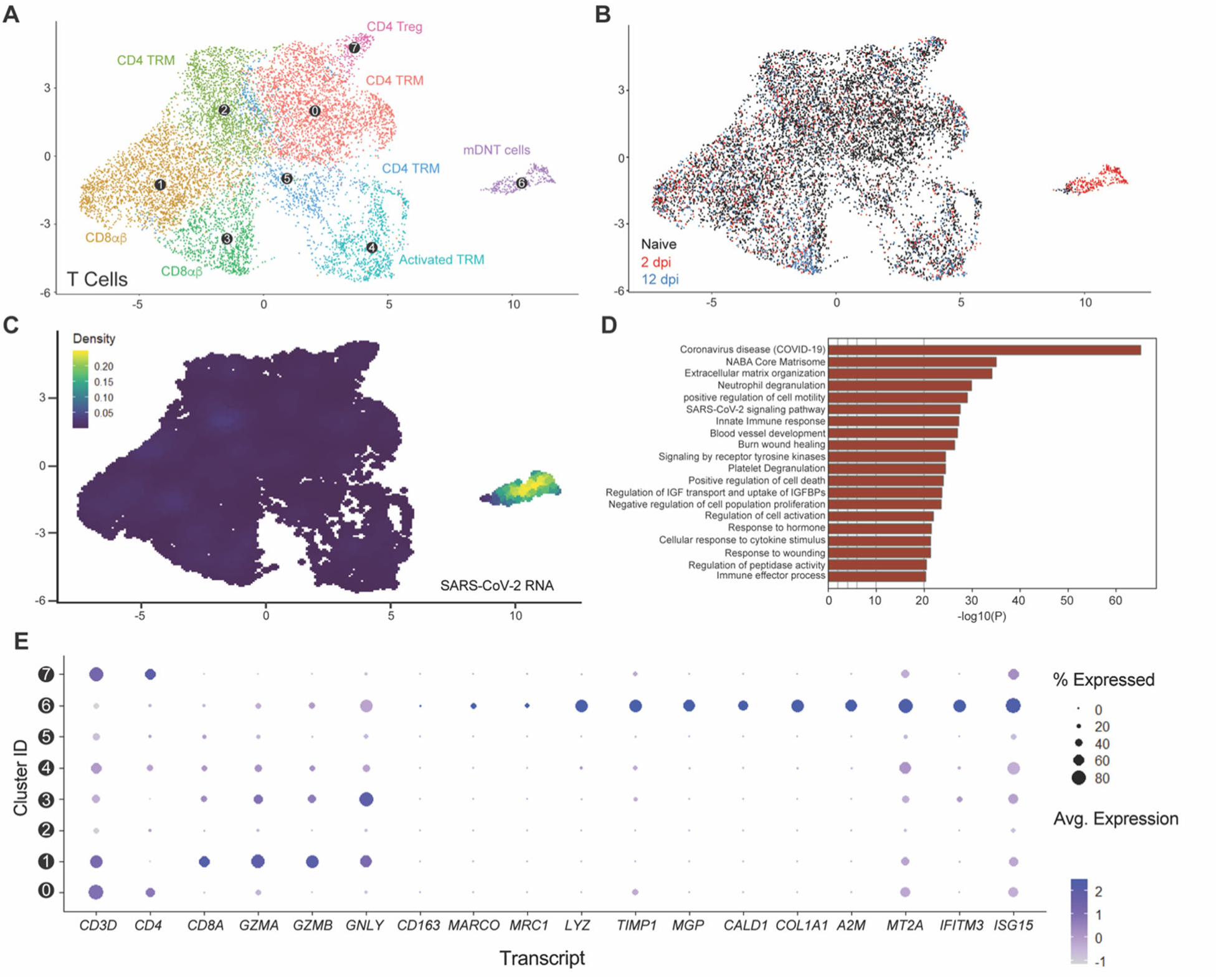
Myeloid-like double-negative T-cell emerges upon acute infection and are enriched in viral RNA. **(A)** UMAP plot showing the sub-clustering of the human T-cell compartment of fLX across all time points. **(B)** Temporal annotation of the human T-cell sub-clusters on the UMAP plot: naïve (black), 2 dpi (red),2 dpi (blue). **(C)** UMAP plot (all time points) showing the distribution of SARS-CoV-2 viral RNA transcripts in the T-cell compartment. **(D)** Go-term analysis showing the most highly upregulated signaling pathways in the mDNT (sub-cluster 6). **(E)** Relative expression of T cells, macrophage, mesenchymal and antiviral markers within each of the T-cell sub-clusters (From 0 to 7; as labeled in panel A).

### Transient antiviral responses by a viral RNA-enriched inflammatory monocyte subset define lung myeloid responses driving SARS-CoV-2 infection resolution

Sub-clustering of myeloid lineages unveiled diverse subpopulations, including alveolar and macrophages (AM) and IM, various monocyte subsets, and one conventional DC subset (cDC) (**Fig. 7A and Fig. S5A-C**). Across all time points, monocytes represented the largest sublineage, which was divided into three subgroups: naïve/resting, patrolling intravascular monocytes (PIM), and inflammatory monocytes (iMO). iMO and AM were the only two subclusters increasing in frequencies upon acute infection at the relative expense of cDC and IM (**Fig.7B,C**). An increase in frequencies within these subclusters was associated with the emergence of distinctive iMO and AM cell populations with infection-induced transcriptomic signatures, underscoring a direct response of these subclusters against viral infection. Following infection resolution, most iMO had phased out and frequencies of alveolar macrophages were concomitantly reduced. Conjointly, the relative number of IM, cDCs and PIM increased through the recruitment of cell populations with distinctive transcriptomic identities from naïve and infection-associated cell populations (**Fig. 7B**, blue).

**Figure 7.**
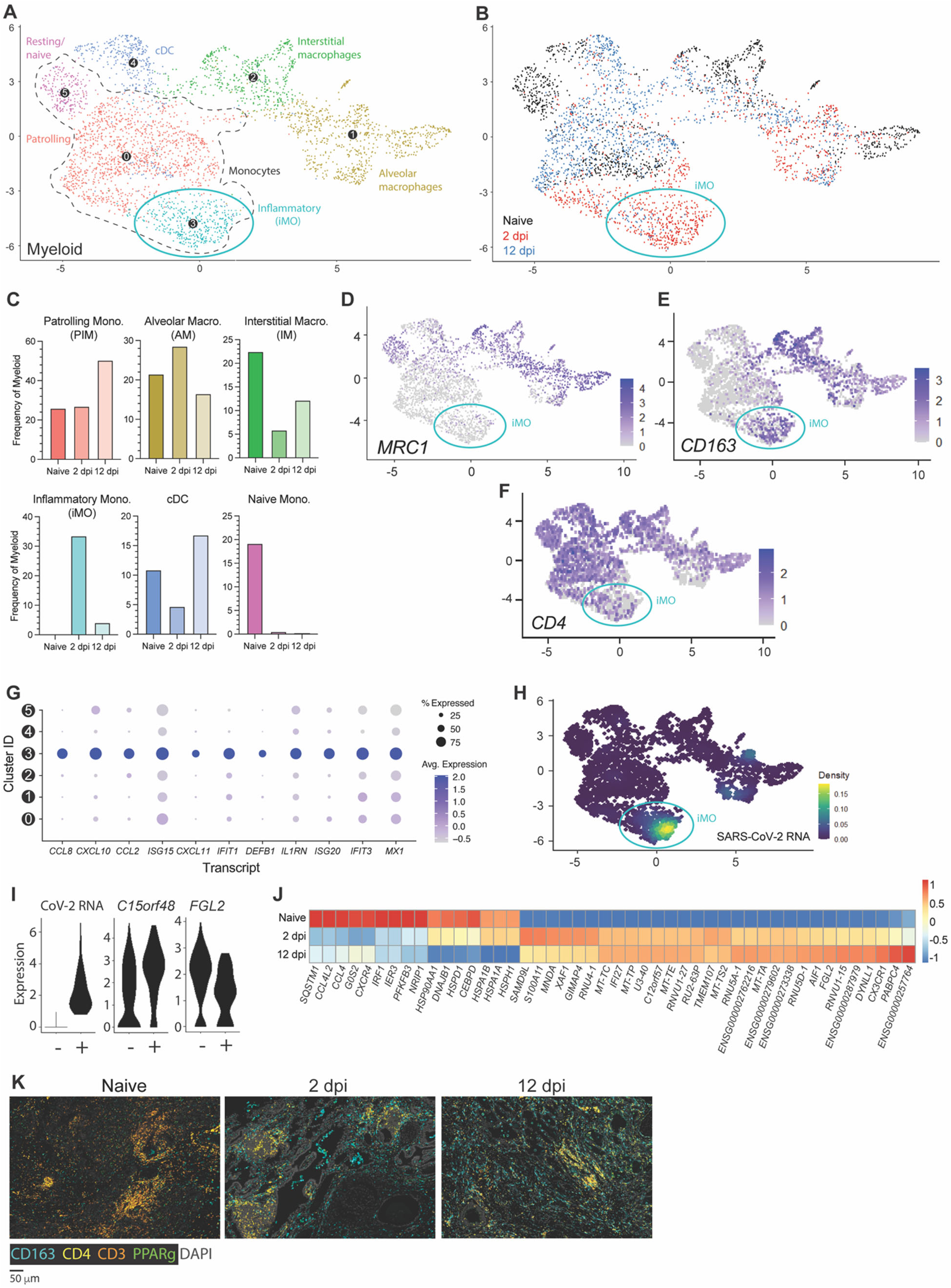
CD163+ extravascular inflammatory monocytes arise during acute viral infection and are predominant antiviral mediators. **(A)** UMAP plot showing sub-clustering of the human myeloid compartment of fLX across all time points. **(B)** Temporal annotation of the myeloid sub-clusters on the UMAP plot: naïve (black), 2 dpi (red), 12 dpi (blue). iMO sub-cluster is indicated with a blue circle. **(C)** Frequency of each myeloid sub-cluster by timepoint. **(D-F)** UMAP plot showing the expression of *MRC1* (D), CD*163* (E), and *CD4* (F) across myeloid sub-sets. **(G)** Relative expression of highly upregulated ISGs and inflammatory markers in each of the myeloid sub-clusters. **(H)** UMAP plot (all time points) showing the distribution of SARS-CoV-2 viral RNA transcripts in the myeloid compartment. **(I)** Violin plots showing expression level of differentially expressed genes between SARS-CoV-2 positive and negative ExiMO (sub-cluster 3). **(J)** Heatmap displaying the relative expression (from -1 to 1) of the top differentially regulated genes over the course of infection within the patrolling monocyte (0) compartment. **(K)** Multiplex fluorescent immunohistochemistry of naïve fLX and fLX at 2 and 12 dpi. CD163: teal, CD4: yellow, CD3: orange, PPARγ: green, Dapi: gray. Two representative images. Scale bar = 100 μM.

Despite the moderate expansion of AM at 2 dpi, iMO were the most abundant myeloid lineage at 2 dpi (33.5%; **Fig. 7B, C**) and uniquely defined among other monocyte subclusters through elevated expression of *VCAN, S100A8, CD14, CD163* and absence/minimal expression of *CD4, MARCO and CD206 (MRC1)* (**Fig. 7D-F and Fig. S5D)**. iMO were also the leading mediators of antiviral responses across all other myeloid subclusters, as exemplified by the robust upregulated expression of interferon-stimulated genes (ISGs) and inflammatory cytokines (*CCL8, CXCL10, CCL2, ISG15, DEFB1, IL1RN, IFIT1, ISG20, IFIT3,* and *MX1*), as well as inflammatory markers such as *CD163* (**Fig. 7E,G**). While some ISGs (e.g., *ISG15, IFIT3, and MX1*) were expressed in other myeloid lineages at 2 dpi, their expression was markedly lower compared to iMO. Notably, upregulation of *CCL8*, *CXCL11* and *DEFB1* transcripts were the most exclusive to iMO (**Fig. 7G**). iMO were also highly enriched in viral RNA, and corresponded to the previously referred 2 dpi-specific viral RNA-enriched subcluster B (**Fig. 3B, 4B, and 7A,B,H**), which also suggested an association between enrichment in viral RNA and potentiated antiviral responses. Concomitantly, we examined whether viral RNA-associated iMOs (CoV-iMOs) displayed a specific transcriptomic signature compared to iMOs (noCoV-iMOs) with undetectable viral RNA. Notably, only two genes correlated with the presence of viral RNA in iMOs: *FGL2 and C15orf48* (**Fig. 7I**). While *FGL2* was down-regulated in CoV-iMO compared to noCoV-iMOs, *C15orf48* was upregulated (**Fig. 7I**). Soluble FGL2 exerts immunosuppressive functions (notably by inhibiting the NF-kB pathway) (*40*). Conversely, the mitochondrial protein C15orf48 is positively regulated by NF-kB signaling (*41*) and has previously been implicated in severe COVID-19, acting as a positive regulator of inflammation (*42*). A recurring feature of lung monocytes in fatal cases of COVID-19 is the expression of *IL-1β* (*1, 2, 6-8*), and inflammasome activation has been associated with the non-productive infection of monocytes(*27*). However, no *IL-1β* expression or inflammasome activation was detected in CoV-iMOs despite enrichment in viral RNA.

In contrast to iMO, *CD4+ CD163-CD206(MRC1)-*PIM were detectable at all time points (**Sub-cluster 0**; **Fig. 7 A-C**). 2 dpi-specific PIM harbored a distinctive, intermediate transcriptomic signature bridging naïve PIM and 2 dpi-specific iMO, and that was defined by the upregulation of specific interferon-stimulated genes (ISGs) such as *IFI27 and XAF1*, and pro-inflammatory genes (*S100A11*) (**Fig. 7J)**. At 12 dpi, PIM was the dominant subset over other monocyte subsets (50.3%; **Fig. 7C**) and displayed a unique transcriptomic profile associated with anti-inflammation, tissue repair and cellular debris clearance mechanisms, notably through the upregulation of *FGL2, DYNLL1* and *CX3CR1*. At 12 dpi, expanded cDC and IM populations also elicited comparable transcriptomic signatures.

mIHC analysis supported our scRNA seq findings. We observed a significant increase in CD4- CD3- CD163+ cells in fLX in the extravascular interstitium and alveolar spaces, mirroring the 2 dpi-specific expansion of CD4- CD163+ iMO and alveolar macrophages uncovered through scRNA seq (**Fig. 7K**). CD3- CD4+ CD163- at that time point could be associated with the 2 dpi-specific PIM population infiltrating the infected fLX. mIHC also recapitulated the reduction of CD3+ T-cells (**Fig. 7K**) at 2 dpi. At 12 dpi, fLX were still enriched in CD4+ cells, likely reflecting PIM infiltration (**Fig. 7K**). CD163+ cells also persisted in fLX at 12 dpi, which can be explained by the combined expansion of IM (CD4+ CD163+) and presence of AM (CD4- CD163+) at that time point as CD163+ iMO phase out (**Fig. 7K**).

Collectively, our findings underscore a coordinated myeloid mobilization to infection resolution and tissue repair, with iMO concentrating viral materials and robust antiviral responses. As the viral materials are resolved, iMO populations dissipate, opening niches for other myeloid lineages to engage in a coordinated tissue repair process.

### Mesenchymal and endothelial signatures of infection resolution

We then examined the contribution of the mesenchymal and endothelial compartment in mediating effective infection resolution. Relative expansion of the mesenchymal compartment upon infection was mainly driven by an increase in fibroblast populations (**Fig. 3C**; **Fig. 8A,B**). An increase in alveolar fibroblasts was observed at 12 dpi (49.6%; naïve=28.1%), which was preceded by the emergence of a 2 dpi-specific cluster of activated fibroblasts (26.5%; naïve=8.9%) (**Fig. 8A-C**), consistent with effective tissue repair in response to lung damage. Activated fibroblasts were the dominant mesenchymal population in mediating antiviral responses (**Fig. 8D**), as displayed by robust upregulation of major pro-inflammatory mediators such as *CXCL10* and *TIMP1* (**Fig. 8F**).

**Figure 8.**
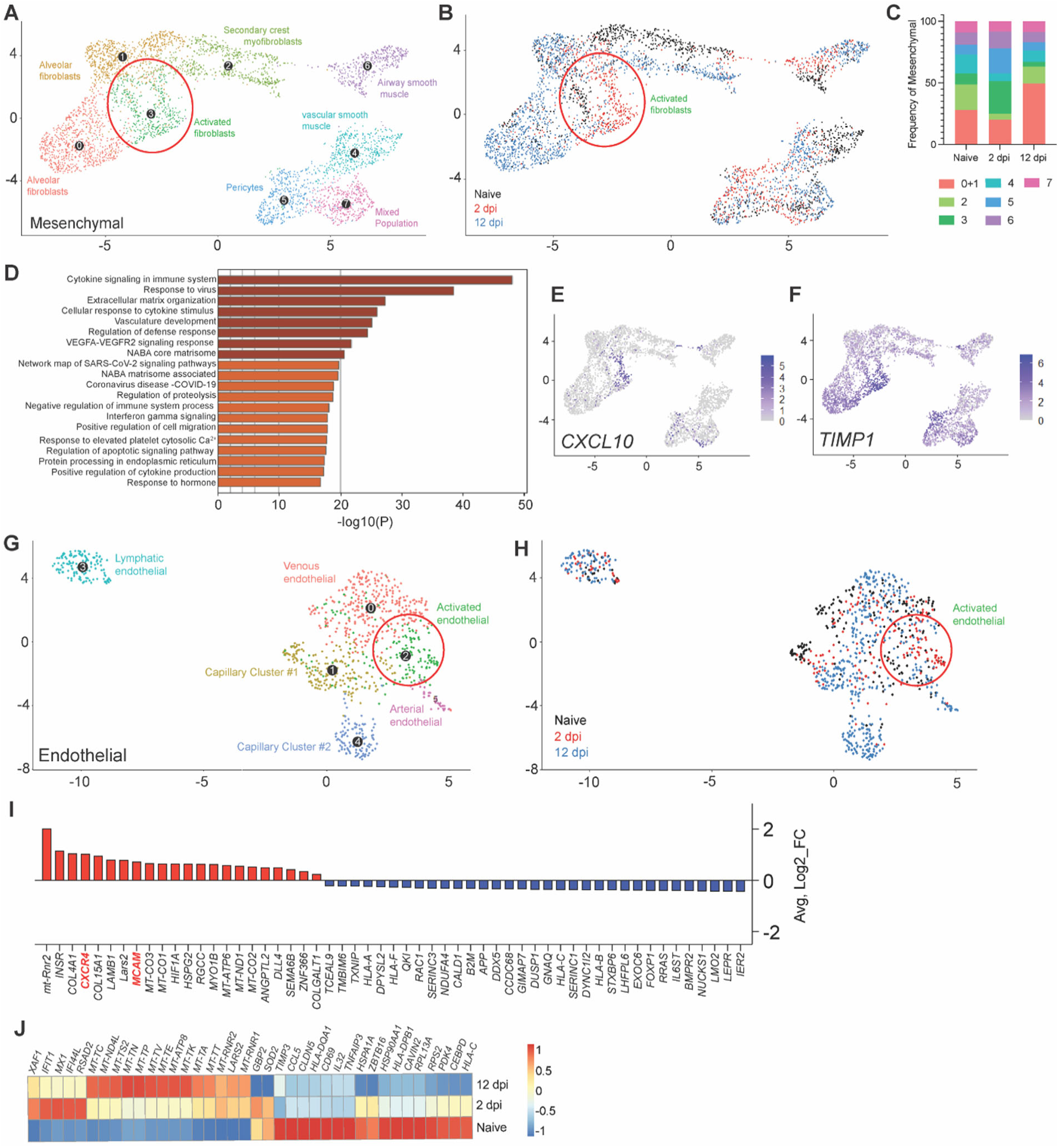
**(A)** UMAP plot sub-clustering of the human mesenchymal compartment in fLX across all time points. **(B)** Temporal annotation of the mesenchymal sub-clusters on the UMAP plot: naïve (black), 2 dpi (red), 12 dpi (blue). **(C)** Frequency of mesenchymal sub-clusters. **(D)** Go-term analysis showing cellular pathways linked to significantly upregulated genes within the activated fibroblasts sub-cluster. **(E-F)** UMAP plot showing *CXCL10* **(E)** and *TIMP1* **(F)** expression within all human mesenchymal sub-clusters and across all time points. Activated fibroblasts are indicated by a red circle in A and B. **(G)** UMAP plot sub-clustering of the human endothelial compartment of fLX across all time points. **(H)** Temporal annotation of the endothelial sub-clusters on the UMAP plot: naïve (black), 2 dpi (red), 12 dpi (blue). **(I)** List of upregulated (red) and downregulated (blue) genes specific to the activated endothelial sub-cluster (Cluster 2) over the other endothelial sub-clusters (*p.adj*≤0.05). **(J)** Heatmap displaying the relative expression (from -1 to 1) of the top differentially regulated genes over the course of infection within the venous endothelial compartment.

The endothelial compartment is an important modulator of immune recruitment through cytokine/chemokine signaling. Infected fLX showed the emergence of an activated, 2 dpi-specific endothelial cell cluster (**Fig. 8G,H**), which was identified by increased expression of several transcripts involved in myeloid chemotaxis (**Fig. 8I**), including *CXCR4* and *MCAM* (*43, 44*). Notably, venous endothelial cells also displayed upregulation of a panel of interferon-stimulated gene transcripts (*XAF1, IFIT1, MX1, IFI44L, RSAD2)* at 2 dpi (**Fig. 8J**) and the downregulation of several transcripts coding for activation markers (*CD69*), MHC genes (*HLA-DPB1, HLA-C*) and major proteins regulating cellular metabolism and transcription (*HSP90AA1, RPL13A*), which was reflective of endothelial dysfunction and stress (**Fig. 8J**) consistently with human patient reports and animal studies (*45-48*). Collectively, our findings support the contribution of the mesenchymal and endothelium compartments in driving antiviral responses and myeloid chemotaxis, respectively, in driving infection resolution.

### Systemic depletion of CD4+ cells abrogates viral clearance in fLX

Many of our findings underscore a robust association between monocyte recruitment into fLX and SARS-CoV-2 infection resolution. This includes: 1) the recruitment of CD4+ PIM into infected fLX, 2) the monocyte nature of iMO and of their dominant antiviral responses, 3) the high enrichment in viral RNA of iMO during infection and 4) the endothelial-mediated myeloid chemotaxis signature at 2 dpi. These pieces of evidence also complement recent human findings which associate effective control of SARS-CoV-2 infection in the nasopharynx with monocyte recruitment(*14*). Antibody-mediated depletions are commonly used to deplete specific hematopoietic subsets, and they are particularly amenable to HIS mouse model studies; further emphasizing the power of these *in vivo* platforms to mechanistically dissect immunological mechanisms in a human context. However, no anti-human CD14 antibodies have been well characterized for *in vivo* depletion of human monocytes. In contrast, anti-human CD4 antibodies have been. To experimentally validate the importance of the recruitment of circulating monocytes to drive infection resolution in fLX, we therefore performed systemic depletion of CD3+, CD4+, and CD8+ cells through the administration of OKT3, OKT4, or OKT8 depleting antibodies, respectively, via intraperitoneal injection of BLTL mice both prior to and after infection (**Fig. 9A**). Human monocytes, as well as some macrophages and dendritic cells (DC) express CD4 (*49, 50*), and the use of these three depleting antibodies will allow us to deconvolute the distinctive impact of T-cells and monocytes in driving infection resolution.

**Figure 9.**
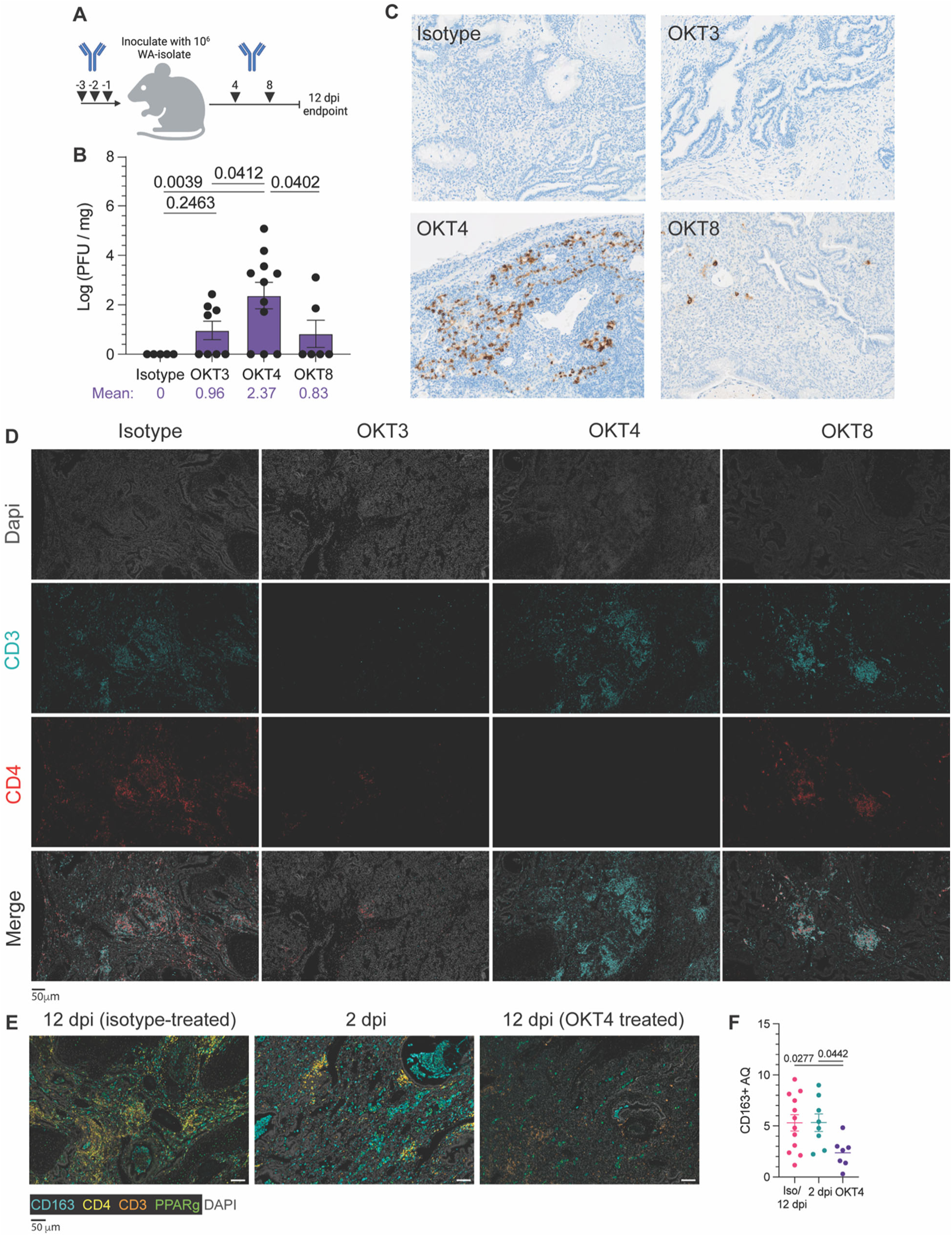
Systemic depletion of CD4+ cells results in persistent infection of fLX. **(A)** BLT-L mice were administered 200 μg of anti-CD3e (OKT3), anti-CD4 (OKT4), anti-CD8 (OKT8) or IgG2a isotype. **(B)** Viral titer (log(PFU/mg)) in fLX extracted from BLT-L mice treated with isotype, OKT3, OKT4, or OKT8 antibody at 12 dpi. **(C)** Immunohistochemistry for SARS-CoV-2 N protein on fLX of depleted or non-depleted BLT-L mice at 12 dpi. **(D)** Multiplex immunohistochemistry on fLX of depleted or non-depleted BLT-L mice at 12 dpi. Dapi = gray, CD3 = teal, CD4 = red. Scale bar = 50 μm. **(E-F)** Multiplex fluorescent immunohistochemistry (E) and CD163+ area quantification (AQ) (F) of fLX that resolved infection (12 dpi, isotype treated), or eliciting acute (2 dpi) or persistent infection (12 dpi, OKT4 treated). CD163: teal, CD4: yellow, CD3: orange, PPARγ: green, Dapi: gray. Scale bar = 50 μM.

Flow cytometry confirmed effective systemic depletion in the blood (**Fig. S6A**). All animals were euthanized at 12 dpi to assess for the persistence of SARS-CoV-2 infection in fLX. While depletion of CD3+, CD4+, and CD8+ cells induced persistent infection in some or most of the animals, only CD4+ cell depletion (mean Log PFU/mg tissue = 2.37) resulted in a statistically significant defect in infection resolution compared to all other experimental conditions (isotype, OKT3, and OKT8-treated mice) (**Fig. 9B**). The rate of productively infected fLX was also higher in CD4+ cell-depleted animals, with 72% of fLX (8/11) showing infection at 12 dpi compared to 50% (4/8) and 33% (2/6) in CD3- and CD8-depleted mice, respectively (**Fig. 9B**). Additionally, when examining the average viral titer (Log PFU/mg of tissue) of persistently infected fLX, the ones from CD4+ cell-depleted animals were significantly higher compared to CD3+ cell- depleted fLX (**Fig. S6B**), furthering that CD4+ cells have a more consequential impact on driving infection resolution than CD3+ cells. Anti-SARS-CoV-2 N IHC confirmed these findings and superior defect of CD4+ cell-depleted animals to resolve infection (**Fig. 9C**). Using multiplex fluorescent IHC (mIHC), we also validated the reduction in CD3+, CD4+ and CD3+ CD4- cells in fLX from CD3-, CD4+ and CD8+ cell-depleted animals respectively, compared to isotype-treated animals (**Fig.9D**). In the fLX of CD4+ cell-depleted mice, CD4+ cells depletion also associated with significant reduction of CD163+ cells at 12 dpi, underscoring the association between CD4+ infiltration into fLX and CD163+ cell recruitment and differentiation (**Fig. 9E,F**).

Persistent infection in CD4+ cell-depleted animals was also associated with significant downregulation of MHC class I (**Fig. 10**), a phenomenon we similarly observed in acutely infected fLX (2 dpi) (**Fig. 10)** and that has been previously reported in cells with active SARS-CoV-2 replication (*51, 52*). This further emphasizes that depletion of CD4+ cells is associated with defective viral clearance mechanisms. These findings suggest that circulating CD4-expressing cells significantly mediate SARS-CoV-2 infection resolution in BLT-L mice.

**Figure 10.**
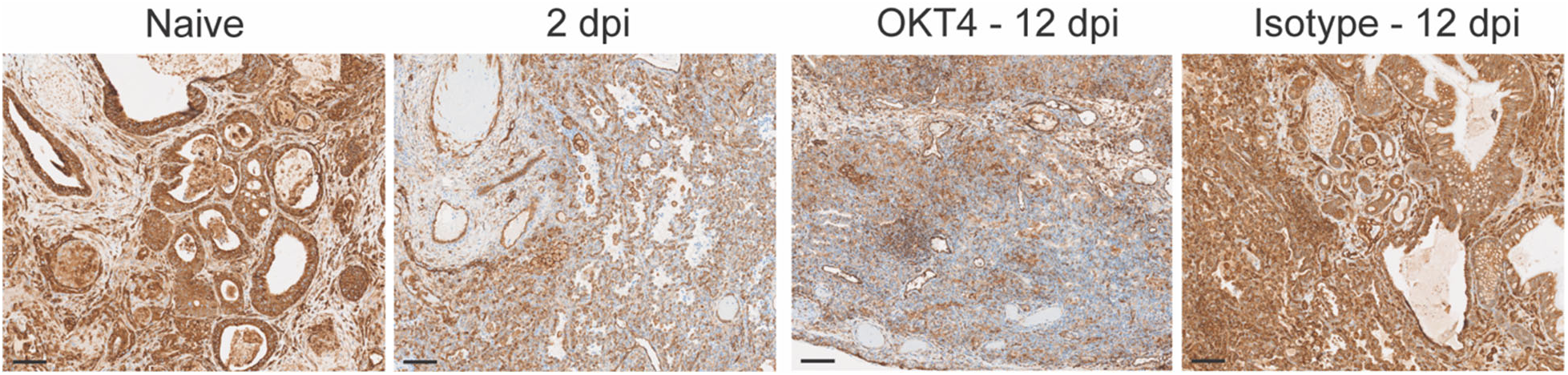
MHC-I staining of fLX tissue sections (anti-MHC class I (EMR8-5) CST 88274) extracted from naïve mice, OKT4-treated mice (12 dpi), or from infected fLX at 2 and 12 dpi. Scale bar = 100 μM

### Defective monocyte recruitment is associated with systemic and local signatures of chronic infection

To further interrogate the impact of CD4+ depletion and monocyte recruitment on SARS-CoV-2 infection, we investigated fLX antiviral responses during acute and persistent infection by quantifying the concentration of 32 cytokines in the peripheral blood of naïve, acutely infected, persistently infected or recovered BLTL mice. Systemic levels of human CCL2 and CCL3, major monocyte attractants, were elevated in CD4+ cell-depleted mice and comparable to those of acutely infected mice (**Fig. 11A,B**). The maintenance of myeloid chemotaxis signals in persistently infected fLX further underlines the contribution of myeloid recruitment in SARS-CoV-2 infection resolution. Notably, among all cytokines and chemokines analyzed, CXCL10 was the only one displaying significantly increased serum levels in acutely infected mice (2 dpi) prior to returning to undetectable levels upon infection resolution (12 dpi) (**Fig 11C**). Several human subsets within fLX express *CXCL10* upon acute infection, including myeloid, mesenchymal and endothelial subsets (**Fig. 11D,E**). However, the myeloid compartment was the major source of *CXCL10* among all human clusters at 2 dpi (**Fig. 11D,E**) and this phenotype was dominantly driven by iMO (**Fig. 7G and 11E**). Therefore, our findings suggest an association between circulating human CXCL10 during acute infection and the simultaneous differentiation of iMO. Notably, levels of circulating human CXCL10 in persistently infected CD4+ cell-depleted mice were not statistically different than those of mice that resolved infection (12 dpi) (**Fig. 11C**). These results emphasize a link between persistent infection and lack of effective CXCL10 responses, further strengthening a connection between monocyte recruitment, iMO antiviral responses and SARS-CoV-2 infection resolution.

**Figure 11.**
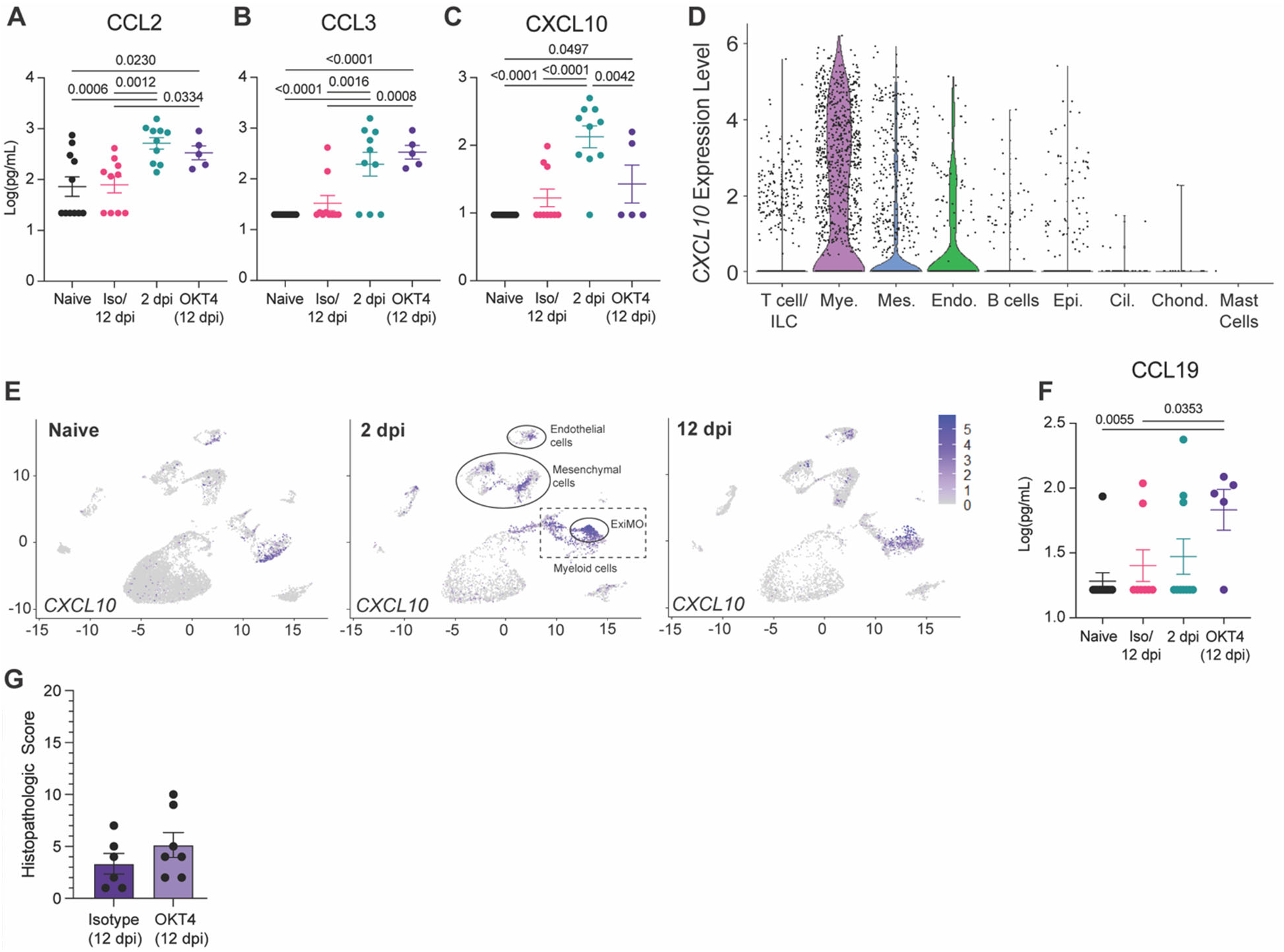
CD4+ cell depletion results in signatures of chronic infection. **(A-C)** Cytokine quantification (**A**: CCL2, **B**: CCL3, **C**: CXCL10) in the serum of naïve, infected (2 dpi and 12 dpi isotype-treated or not) and OKT4-treated BLT-L mice (12 dpi). **(D)** Violin plot showing expression level of *CXCL10* per cell and within each human lineage. **(E)** UMAP plot showing *CXCL10* expression within all human lineages in naive fLX and at 2- and 12 dpi. Locations of the mesenchymal, endothelial, myeloid and ExiMO clusters at 2 dpi are indicated. **(F)** Quantification of CCL19 in the serum of naïve, infected (2 dpi and 12 dpi isotype-treated or not) and OKT4- treated BLT-L mice (12 dpi). **(G)** Histopathological scoring of fLX extracted from isotype and OKT4-depleted BLT-L mice (12 dpi). Error bars indicate mean ± Standard error of the mean. One*- way ANOVA, t-test. p-values are indicated on graphs*.

Persistently-infected mice antiviral responses were also distinguishable from acutely infected (2 dpi) and isotype-treated (12 dpi) mice by elevated levels of CCL19 in serum (**Fig. 11F**). CCL19 is a pro-inflammatory cytokine that has been linked with persistent viral replication and inflammation such as in the context of HIV-1 infection (*53*), underlining that persistent SARS-CoV-2 infection in BLT-L mice recapitulate key immunological features of chronic viral infection.

We have previously reported that fLX exhibit significant histopathological manifestations of disease upon SARS-CoV-2 infection in the absence of an engrafted human immune system (3), highlighting that tissue damage in fLX is virally induced. However, persistent infection did not result in any significant histopathological manifestations of disease in fLX, which strongly contrasts with acutely infected fLX (**Fig. 2H**, **Fig. 11G**). As minimal tissue damage is a hallmark of chronic viral infection (*54*), these findings further support evidence that CD4+ cell depletion promotes tissue remodeling processes underlying chronic infection and lasting viral persistence.

## DISCUSSION

As our appreciation of the immunological differences between mice and humans continues to grow, humanized mouse models increasingly stand out as robust platforms to understand how viral pathogens interact with human tissues and the human immune system. These models are especially valuable when investigating tissue and mucosal immunity since such investigations remain impractical in human patients.

The SARS-CoV-2 pandemic has emphasized the need to increase our understanding of immune mechanisms that can drive protection against immunologically novel respiratory viruses. Mice engrafted with human immune systems and human lung tissues have emerged as valuable tools for such investigations (*21, 23-25*), bridging the limitations of conventional animal models and the challenges associated with human studies. We previously reported using the HNFL mouse model that such models can be leveraged to capture immunological signatures defining effective control of SARS-CoV-2 infection (*23*). However, the enhanced myeloid reconstitution of the HNFL model rapidly inhibits viral replication in fLX, precluding our ability to study protective naive immunological mechanisms at play upon acute and potentially symptomatic infection. This study aimed to uncover novel facets of these mechanisms using a humanized mouse model that exhibits robust susceptibility to acute SARS-CoV-2 infection prior resolving infection in a hematopoietic-dependent manner.

We found that BTL-L mice are able to effectively clear infectious viral particles following an early peak of viral replication in fLX. Infection resolution was associated with rapid mobilization of the human immune cells into fLX upon viral inoculation, which aligns with previous evidence that immunodeficient mice only engrafted with fLX are unable to clear infection (*22, 23*). Using scRNAseq analysis, we identified a comprehensive network of novel factors involved in the resolution of SARS-CoV-2 infection (**Fig. 12**). Our study identified three major hallmarks of this process: 1) The recruitment of iMO, which display high level of enrichment in viral RNA and mount dominant antiviral responses across all myeloid lineages, notably defined by CXCL10 expression; 2) The differentiation of mDNT cells, which are highly enriched in viral RNA and exhibit non-canonical macrophage features and cytotoxic signatures; And 3) The synergistic contribution of endothelial and mesenchymal cells in infection resolution via potentiating antiviral responses and myeloid chemotaxis, respectively. Consistent with monocyte infiltration into infected fLX being a dominant feature of our infection resolution model, systemic depletion of CD4+ cells, but not T-cells, abrogated viral clearance. Persistent infection is associated with the lack of systemic CXCL10 responses, dominantly mediated by iMO, as well as with signatures of chronic infection, including systemic CCL19 expression and lack of fLX damage.

**Figure 12.**
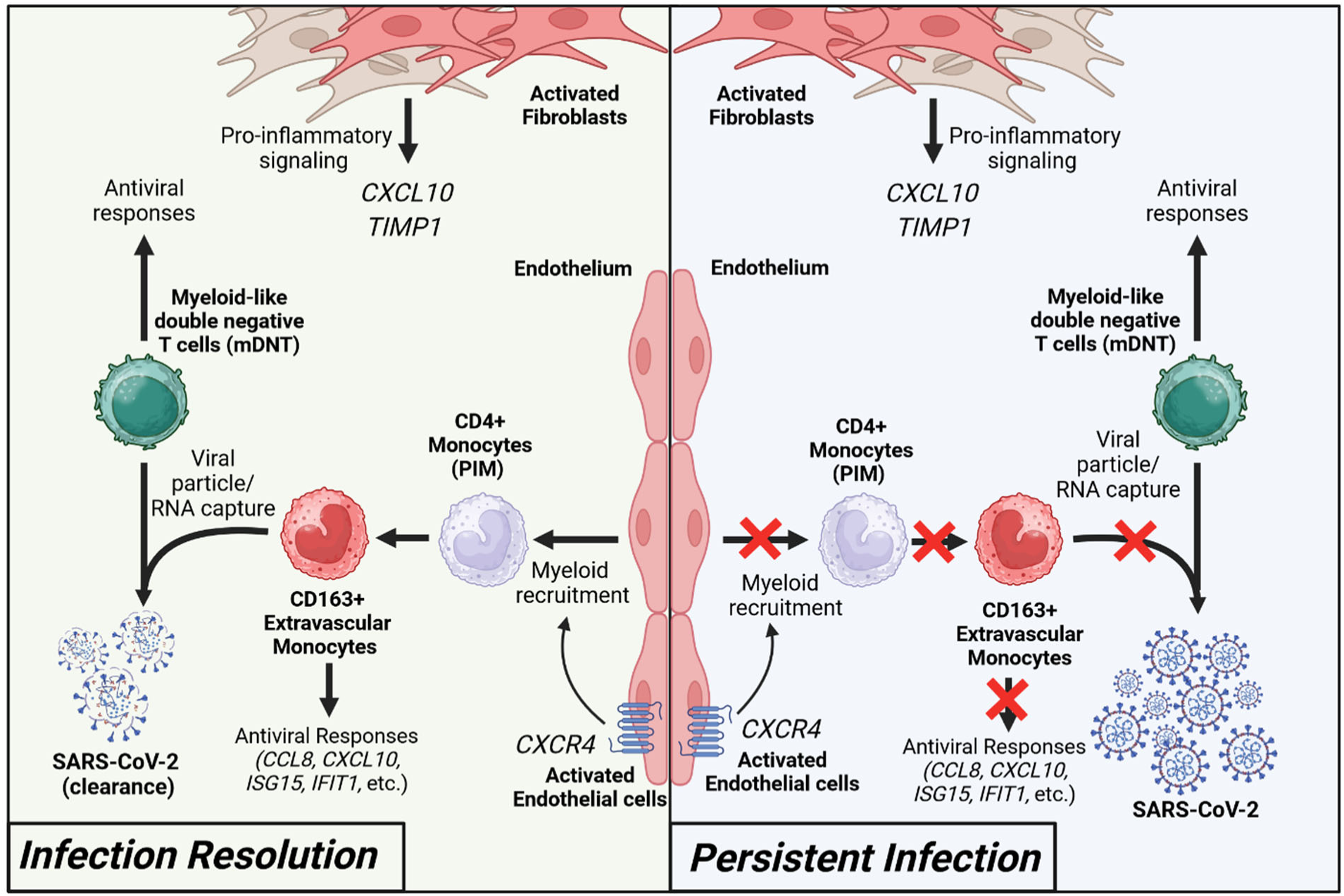
Immune signatures of SARS-CoV-2 infection resolution in human lung tissues. During effective viral resolution and tissue repair (left panel), myeloid cells are recruited to the site of infection and differentiates into CD163+ inflammatory monocytes (iMO). iMO produce antiviral and inflammatory signals and capture viral RNA/particles. Additionally, myeloid-like double negative T cells (mDNT), act to enhance the capture of viral RNA/particles and antiviral responses. In parallel, activated endothelial cells and fibroblasts contribute to the recruitment of myeloid cells and inflammatory response, respectively. Collectively, these events lead to viral clearance and infection resolution. In the context of persistent infection (left panel), absence of effective CD4+ monocyte recruitment prevents infection resolution.

Our findings empower a recent human study reporting that protection from SARS-CoV-2 infection is associated with a rapid monocyte response in the nasal cavity and a decreased number of circulating monocytes(*14*). Specifically, our work extends the findings of this human study beyond the limitations of the human model by exploring tissue-resident events, unraveling the identity of monocyte populations that extravasate tissues, differentiate and mount tissue-resident immune responses to clear infection. By providing enhanced resolution on key protective immunological processes and mediators that human studies alone cannot capture, our works underscores how human and humanized mouse studies can effectively complement themselves for improving our understanding of human antiviral immunity.

To the best of our knowledge, this study represents the initial evidence of a direct role played by human lung extravascular inflammatory monocytes in the resolution of respiratory viral infections. Extravascular monocytes have been proposed to serve as immune sentinels through their position at the interface of the lung capillaries and alveoli (*55, 56*). However, although previous research reported that these cells can promote T-cell resident memory differentiation following viral infection (*57*), their direct antiviral functions have not been documented until now. Four major features characterized these cells: 1) a dominant CD163-expressing population that emerges during acute infection before dissipating, 2) a major source of CXCL10 expression, 3) a high enrichment in viral RNA, and 4) the induction of robust antiviral responses.

Several lines of evidence support the monocyte nature of these cells. They display a close transcriptomic relationship with patrolling monocytes, which have been reported to give rise to transient, non-classical extravascular (including alveolar) monocytes in the mouse lung (*56*). The iMO gene expression profile is also similar to those of human FCN1-monocytes recovered from the broncho-alveolar lavage of COVID-19 patients with acute respiratory disease syndrome (*58*). While further studies are needed, our data collectively suggest CD4+ PIM infiltrating the infected fLX may differentiate into iMO to promote robust antiviral responses. Furthermore, our data show that some iMO are negative for viral RNA, suggesting that such differentiation is independent of an association with viral materials, although the presence of viral RNA could potentiate antiviral responses.

The fate of iMO following infection resolution also remains unclear and will have to be further deciphered. By 12 dpi and infection resolution, fLX display residual iMO and are enriched in CD4+ PIM and CD163+ CD206+ IM. CD206+ IM are involved in response to wounding and infection recovery (*56*) and CD163+ CD206+ monocyte-derived IM (*58*) have been shown to directly derive from extravasating, circulating CD14+ monocytes in another human immune system-engrafted mouse model (*59*). Therefore, one hypothesis would be that the pool of CD163+ CD206+ IM is derived both from iMO and newly recruited circulating monocytes.

Our findings also reveal that different monocytic fates are likely associated with distinct clinical outcomes of SARS-CoV-2 infection. While our study emphasizes the protective role of monocytes in preventing severe COVID-19, monocytic lineages have also been associated with severe COVID-19(*2, 5, 36, 58*). Excessive monocyte infiltration, macrophage inflammation and fibrotic response can lead to potentially fatal acute respiratory distress syndrome (ARDS) and fibrosis despite promoting infection resolution. A suspected driver of excessive myeloid inflammation is the ability of SARS-CoV-2 to trigger abortive infection of both monocytes and macrophages(*27, 60-62*) through viral RNA replication and protein production, which results in inflammasome activation(*27, 28, 63*). Consistently, viral RNA is enriched in inflammatory monocytes and macrophages in lung autopsy samples from fatal COVID-19 cases (*5*). However, our findings that viral RNA in iMO is associated with infection resolution and tissue protection suggest that rapid clearance of viral RNA-enriched inflammatory monocytes, as opposed to their persistence, is associated with favorable clinical outcomes.

Lung extravascular monocytes deriving from PIM also exhibit a dynamic transition state and can differentiate into CD206-IM(*56*), primary responders to infection and important drivers of inflammation. However, no CD206- macrophages were observed in infected fLX at any time point. The transient presence of viral RNA-enriched iMO in fLX, leading to the absence of iMO differentiation into CD206- IM, could therefore mitigate the risk of uncontrolled inflammation during infection resolution. This process could prevent severe tissue damage and excessive extravasation of circulating monocytes, which would otherwise differentiate into pro-fibrotic CD163+ CD206+ IM, a major driver of ARDS(*58*).

Understanding the cellular and molecular players defining the fate of iMO upon exposure to SARS-CoV-2 and how key regulatory crossroads in this subset may result in differential clinical outcomes is of particular interest. Compounding immune dysregulations, such as elevated inflammatory baseline and/or epigenetic imprinting related to innate immune training (*64*), may hinder PIM and/or iMO’s ability to adequately regulate their inflammatory responses in a timely manner upon encountering viral materials or inflammatory clues. This could also favor extended cell survival (NF-κB is both involved in inflammasome activation and cell survival (*65*)) and subsequent differentiation into inflammatory macrophages, fostering exacerbated inflammation. Our study also creates a mandate for probing the ubiquitous nature of iMO antiviral responses during other respiratory viral infections and investigating more comprehensively the antiviral functions of this novel subset beyond SARS-CoV-2 infection.

In parallel to ExiMO, we also identified mDNT cells, a transient DNT cell population highly enriched in viral RNA and only observed upon acute infection, and which exhibits expression of key macrophage-defining genes and cytotoxic markers. This population displays similarities with a subpopulation of cytotoxic DNT cells previously reported in mouse spleen(*38*) and with innate-like T-cells(*66*), which also harbor cytotoxic signatures. mDNT also share transcriptomic similarities with γδ T-cell subsets previously described as bearing myeloid and cytotoxic functions(*67, 68*). However, mDNT appear unique through a specific blend of lymphoid, and conventional macrophage and mesenchymal markers and their short-lived nature (suggested by their complete absence at 12 dpi). While CD8+ T-cells and macrophages isolated from nasopharyngeal swabs have been identified to harbor SARS-CoV-2 RNA in human challenge studies(*14*), mDNT did not meet the canonical transcriptomic signatures of these subsets, suggesting the existence of tissue-resident events driving specific cellular differentiation processes upon infection. Additional investigations are required to better understand the identity, fate and functions of this cell subset in lung antiviral immunity. However, it is tempting to speculate that mDNT may uniquely synergize CD8+ cytotoxic T-cell functions with viral particle phagocytosis to effectively control infection prior to undergoing rapid cell death.

Despite our findings related to mDNT, the contribution of T-cells in SARS-CoV-2 infection resolution remains elusive in our model. CD3+ cell depletion resulted in a partial abrogation of infection resolution, with 50% of animals still capable of clearing infection. In addition, even when viral infection persisted in CD3+ cell-depleted animals, viral titers in fLX remained significantly lower than in the fLX of persistently infected CD4+ cell-depleted animals. One possible interpretation is that T-cell depletion delays but does not abolish infection resolution. CD8+ cell-depleted mice may display delayed viral clearance compared to isotype-treated mice (with resolution occurring between 6dpi and 12 dpi), and CD3+ T-cell-depleted mice may be undergoing progressive resolution by 12 dpi until complete clearance by a later time. In contrast, the viral titer observed in fLX of CD4+ cell-depleted mice show no evidence of ongoing infection resolution. How CD3+ depletion may impact mDNT differentiation, and how T-cells (including mDNT) and ExiMO functions may synergize to rapidly resolve infection without extensive inflammation-induced tissue damage will need further investigation.

There are inherent limitations associated with the BLT-L mouse model. First, direct viral inoculation into fLX potentially bypasses key immune checkpoints in the upper respiratory tract, which are not recapitulated in our model. Second, the hematopoietic reconstitution and functions of the BLT-L mouse model remain imperfect, notably through the underrepresentation of specific hematopoietic subsets, including dendritic cells and granulocytes, and limited B-cell responses – which may ultimately bias the dynamics of viral clearance. Despite these limitations, BLT-L mice recapitulated human lung responses to SARS-CoV-2 infection through AT2 loss of programming, fibroblast and endothelial activation, and effective clearance of infection through robust hematopoietic responses. This mouse model also supported viral adaptation processes with public health relevance, and recapitulated signatures of persistent viral replication observed in patients suffering from chronic infection. Our work demonstrates the potential of the BLT-L mouse model to uncover naïve immune mechanisms and mediators governing the effective resolution of lung infection by SARS-CoV-2 and open avenues for a comprehensive examination of such processes against other viral respiratory infections, which may pave the way toward innovative immunotherapy strategies against these diseases.

## AUTHOR CONTRIBUTIONS

D.K., A.B.B. and F.D. conceptualized the overall study. C.H. conceptualized the computational component of this study and oversaw the analysis of the single-cell RNA sequencing data. D.K., A.B.B., N.A.C. and F.D. designed the experiments. D.K., A.K.O., M.S, A.N, H.P.G., M.E., V.V., A.B.B. and F.D. performed experiments. D.K., A.K.O., A.T., J.T., M.S, A.N, P.M., M.E., J.H.C., V.V., N.A.C., C.H., A.B.B. and F.D. analyzed the data. J.T and C.H. carried out computational analysis. M.E. carried out electron microscopy analysis. A.B.B. and V.V. provided access to key resources. D.K. and F.D. wrote the manuscript with contributions from all authors.

## ACKNOWLEDGMENTS

This work was supported in part by a start-up fund and Peter Paul Career Development Professorship from Boston University (to F.D.), grants from the National Institutes of Health (K22 AI144050 to F.D.), Clinical and Translational Science Awards (grant UL1 TR001430) from the National Center for Advancing Translational Sciences of the National Institutes of Health (to F.D., and N.A.C.) and an Evergrande MassCPR award (to NEIDL). A.B.B. was supported by R01AI174875, R01AI174276, DP2DA040254, as well as CDC subcontract 200-2016-91773-T.O.2. and a Massachusetts Consortium on Pathogenesis Readiness (MassCPR) grant. We also thank the Evans Center for Interdisciplinary Biomedical Research at Boston University Chobanian and Avedisian School of Medicine for their support of the Affinity Research Collaborative on ‘Respiratory Viruses: A Focus on COVID19’. This work utilized a Ventana Discovery Ultra and Akoya PhenoImager HT that were purchased with funding from National Institutes of Health SIG grants (S10 OD026983 & S10OD030269). We thank Ronald B. Corley, NEIDL director at the time of this study and MassCPR award recipient, for his constant encouragement and support for this study. We thank the Boston University Animal Science Center, the Ragon Institute Human Immune System Mouse core, the single cell sequencing core and microarray and sequencing core, and the flow cytometry core at Boston University Chobanian and Avedisian School of Medicine, and all the NEIDL animal core staff for their outstanding support. We also thank all the Douam, Balazs, Connor, and Crossland lab members, NEIDL members, and members of the Department of Virology, Immunology, and Microbiology and Pathology at Boston University for their constant support and advice. D.K. is supported by a T32 training grant in immunology (T32AI007309).

## MATERIALS AND METHODS

Detailed descriptions of the materials and methods used for this study are in the Supplemental Materials.

### Institutional approvals

All experiments in this study, including those conducted in BSL-3, were approved by an institutional biosafety committee. Animal experiments described in this study were performed in accordance with protocols that were reviewed and approved by the Institutional Animal Care and Use and Committee of the Ragon Institution and Boston University. All mice were maintained in facilities accredited by the Association for the Assessment and Accreditation of Laboratory Animal Care (AAALAC). All replication-competent SARS-CoV-2 experiments were performed in a biosafety level 3 laboratory (BSL-3) at the Boston University National Emerging Infectious Diseases Laboratories (NEIDL).

### Mouse strains and sex as biological variable

Female NOD.Cg.-*Prkdc^Scid^Il2rg^tm1Wjl^*/SzJ (NSG) mice were obtained from the Jackson Laboratory, catalog number 005557. NSG mice were maintained by the Ragon Institute Human Immune System Mouse core prior to engraftment and shipment to the NEIDL, Boston University. In our study, only female mice were engrafted with human fetal tissues because of their ability to support higher levels of engraftment than males. As we investigated human tissue responses to infection and only leveraged mice as xenorecipients, the sex of the animals does represent a critical variable for our study.

### Human fetal tissues

De-identified human fetal tissues were procured from Advanced Bioscience Resources (Alameda, CA, USA).

### Generation of BLT-L mice

BLT mice were generated via irradiation of female NOD.Cg.- *Prkdc^Scid^Il2rg^tm1Wjl^*/SzJ mice (NSG mice; Jackson Laboratory #005557) prior to implantation of human fetal thymic and fetal liver tissue (Advanced Bioscience Resources) under the murine kidney capsula. Two pieces of homologous human fetal lung tissue were implanted into subcutaneous dorsal pockets of mice. Post-implantation, mice intravenously received 1×10^5^ homologous CD34+ cells. Human immune reconstitution was determined by flow cytometry at weeks post-implantation. Three distinct human donors were used for this study.

### Mouse inoculation with SARS-CoV-2

BLT-L mice were anesthetized with 1–3% isoflurane prior to inoculation via subcutaneous, intra-fetal lung xenograft (intra-fLX) injection with 10^6^ plaque forming unites (PFU) of SARS-CoV-2 WA-1 isolate in 50 μL of sterile 1X PBS. Mice were euthanized at 2-, 6-, and 12-days post inoculation.

### In vivo antibody depletion

BLT-L mice were administered 200 mg of anti-CD3e (OKT3) (BioxCell; cat. # BE0001-2), anti-CD4 (OKT4) (BioxCell; cat. # BE0003-2), anti-CD8 (OKT8) (BioxCell; cat. # BE0004-2), or isotype IgG2a (Thermofisher; cat # 02-6200) antibody 3-, 2-, and 1-day prior to inoculation and 4- and 8-days post inoculation with 1×10^6^ PFU SARS-CoV-2 WA-1.

### Quantification and statistical Analysis

For histopathological score and viral load/titer comparisons a Kruskla-Wallis, non-parametric one-way ANOVA with Benjamini, Krieger, and Yekutieli correction for multiple comparisons was applied given the non-continuous nature of the data. For cytokine data, an Ordinary one-way ANOVA with uncorrected Fishers LSD was used as the data was collected from different time points, treatment conditions, and cohorts. A Kruskla-Wallis, non-parametric one-way ANOVA with an uncorrected Dunn’s test was applied for CD163+ Area quantification (AQ) due to the independent comparisons between the samples. All statistical tests and graphical depictions of results were performed using GraphPad Prism version 9.0.1 software (GraphPad Software, La Jolla, CA). For all tests, p≤0.05 was considered statistically significant. Statistical significance on figures and supplemental figures is labeled with p-values and non-significant values are labeled with n.s. or left unlabeled.

### Data availability

The Genome Expression Omnibus (GEO) accession number to access the raw data of our scRNAseq analysis is GSE255200.

