## Supplemental materials for "Immune Signatures of SARS-CoV-2 Infection Resolution in Human Lung Tissues"

**FOR**

**CONTENT:**

**1. Supplemental Materials and Methods**

**2. Supplemental Figures (6 Figures)**

**Figure S1.** BLT-L mice are susceptible to SARS-CoV-2 infection.

**Figure S2.** SARS-CoV-2 undergoes viral adaptation within fLX.

**Figure S3.** Single-cell RNA sequencing analysis of fLX upon SARS-CoV-2 infection.

**Figure S4.** Gene signature of macrophage-like T cells.

**Figure S5.** Gene signature of myeloid and endothelial sub-clusters.

**Figure S6.** Assessment of systemic depletion efficiency and analysis of viral titers in mice with abrogated viral clearance mechanisms.

**3. Supplemental Tables (1 Table)**

**Table S1.** SARS-CoV-2 mutations identified among viral reads isolated from fLX at 2 dpi.

**4. Supplemental References**

### SUPPLEMENTAL MATERIALS AND METHODS

#### *Resources*

***Cell lines.*** VeroE6 cells were grown in Dulbecco's modified Eagle's medium (DMEM) supplemented with 10% heat inactivated fetal bovine serum (Bio-Techne, R&D systems, Minneapolis, MN, USA) and 1% (v/v) Penicillin Streptomycin (Thermo Scientific, Waltham, MA, USA).

***Antibodies.*** The anti-mouse antibody CD45-PE-Dazzle5 clone 30-F11 was purchased from BioLegend (San Diego, CA, USA) and used for flow cytometry. The following anti-human antibodies were used for flow cytometry and were all acquired from BD Biosciences (San Jose, CA, USA): HLA-DR-BV510 clone L243, CD3-BV605 clone UCHT1, CD20-BV650 clone 2H7, CD16 (Fc$\gamma$RIII)-BV117 clone B73.1, CD45-BV785 clone H130, CD8-FITC clone RPA-T8, CD33-PE clone WM53, CD14-PerCP-Cy5.5 clone HCD14, CD45RA-PE-Cy7 clone HI100, CD56-allophycocyanin clone QA17A16, CD4-Alexa Fluor 700 clone SK3. The following anti-human antibodies were used for *in vivo* depletions: From Bio-X Cell (Lebanon, NH, USA): anti-human CD3 clone OKT3, anti-human CD4 clone OKT4, anti-human CD8 clone OKT8; From Thermofisher: mouse IgG2a isotype. The anti-SARS-CoV-2 (2019-nCoV) Spike RBD-mFc clone #57 used as a control for neutralization assays was acquired from Sino Biologicals.

#### *Experimental methods*

***Generation of BLT-L mice.*** BLT mice were generated via irradiation of female NOD.Cg.-*Prkdc*$^{Scid}$*Il2rg*$^{tm1Wjl}$/SzJ mice (NSG mice; Jackson Laboratory #005557) with 200 rads from an X-ray irradiator. Mice then underwent surgery to implant 1mm$^3$ of human fetal thymic and fetal liver tissue (Advanced Bioscience Resources) under the murine kidney capsula. After the blunt resection of the thoracal subcutis, through the same surgical opening, two pieces of homologous human fetal lung tissue into the subcutaneous dorsal area of the 8-15 week old females. Post-implantation, mice intravenously received $1x10^5$ homologous CD34+ cells. Human immune reconstitution was determined by flow cytometry at weeks post implantation. Three distinct human donors were used for this study.

***Generation of SARS-CoV-2 isolate and recombinant SARS-CoV-2 NanoLuc viral stocks.*** All replication-competent SARS-CoV-2 experiments were performed in a BSL-3 facility at the Boston University National Emerging Infectious Diseases Laboratories. The clinical isolate named 2019-nCoV/USA-WA1/2020 strain (NCBI accession number: MN985325) of SARS-CoV-2 was obtained from BEI Resources (Manassas, VA, USA). Recombinant SARS-CoV-2 expressing NanoLuc Luciferase (rSARS-CoV-2 NL) (Xie et al., 2020) was graciously provided by the Laboratory of Pei-Yong Shei. Viral stocks were grown and purified as previously described (reference HNFL manuscript). Briefly, $1x10^7$ Vero E6 cells were plated in a T-175 flask the day prior to virus generation. The next day, cells were infected with virus diluted in 10 mL of Opti-MEM (ThermoFisher Scientific, Waltham, MA, USA) and incubated for 1 h at 37°C to allow for virus adsorption. After 1 h, 15 mL of DMEM containing 10% FBS and 1% penicillin/streptomycin was added to cells and incubated overnight. The next day, media was removed, cells were rinsed with 1X PBS, pH 7.5 (ThermoFisher Scientific) and 25 mL of fresh DMEM containing 2% FBS was added. Cells were assessed for cytopathic effect (CPE), media was collected, filtered with a 0.22 µm filter, and concentrated by sucrose gradient. Concentrated virus was suspended in sterile 1X PBS, pH 7.5, aliquoted and stored at –80C.

***Viral quantification by plaque assay.*** The titer of our viral stocks was determined by plaque assay. Vero E6 cells were seeded into a 12-well plate at a density of $2x10^5$ cells per well. The next day, cells were infected with 10-fold serial dilutions of the viral stocks and incubated for 1 hour at 37°C to allow for viral adsorption. After 1 hour incubation, 1 mL of overlay media (1.2% Avicel (DuPont, Wilmington, DE, USA; RC-581) in DMEM supplemented with 2% FBS and 1%

Pen/Strep) was added to each well. Three days later, the overlay media was removed, and cells were fixed for 1 hour at room temperature with 10% neutral buffered formalin (ThermoFisher Scientific). Formalin was removed after fixation and cells were stained for 30 min at room temperature with 0.1% crystal violet (Sigma-Aldrich) prepared in 10% ethanol/water. Crystal violet was removed, cells were washed with water and plaque forming unites were counted to determine viral titers.

***Generation of single cell suspension from mouse blood.*** Blood processing was performed as previously described (*1*). Briefly, blood was collected via submandibular bleeding or heart puncture and transferred into EDTA capillary collection tubes (Microvette 600 K3E; Sarstedt, Nümbrecht, Germany). Blood was centrifuged at 3,500 RPM for 10 minutes at room temperature, serum was removed, and whole blood was subject to ACK lysing (ThermoFisher Scientific; #A1049201) for 10 minutes at room temperature. Lysing was quenched with 10% (v/v) FBS DMEM media and cells were washed twice with a 1% (v/v) FBS-PBS solution (FACS Buffer) before downstream processing.

***Generation of single cell suspension from fLX.*** Fetal lung xenografts were processed as previously described (*1*). Briefly, lung tissues were placed in a 60 mm dish, minced using a disposable scalpel, then transferred into a 15 mL conical tube containing 3 mL of digestion buffer (HBSS minus Ca2+, Mg2+, and phenol red, 0.5 mg/mL Liberase TM, 1 mg/mL DNase I) and incubated at 37°C for 30 min with agitation every 10 min. After digestion, tissue was passed through a 70 μm strainer on a 50 mL tube, mashed using the plunger of a 3 mL syringe and washed twice with 1 mL FACS buffer. Cell suspension was centrifuged at 300 *x g* for 5 minutes at 4°C, suspended in 1 mL of ACK lysing buffer, and incubated for 2 min at room temperature. After incubation, lysis was quenched with 12 mL of FACS buffer, cell suspension was centrifuged at 300 *x g* for 5 min at 4°C, and the cell pellet was suspended in 1 mL of FACS buffer prior to downstream processing.

***Viral RNA extraction from serum.*** Viral RNA was extracted from serum using a Zymo Viral RNA extraction kit (Zymo Research, Irvine, CA, USA: #R1035) following the manufacturers protocol. Briefly, serum was mixed with RNA/DNA shield (Zymo) at a 1:1 ratio. RNA buffer was then added to the serum (2:1 ratio) and passed through a column by centrifugation at 13,000 x g. The column was then washed twice, and RNA was eluted with 15 µL of RNase/DNase free water.

***RNA extraction from fLX.*** During necropsy a piece of fLX tissue was placed in 600 µL of RNAlater (MilliporeSigma: #R0901500ML) and stored at −80°C until analysis. For RNA extraction, 20–30 mg of tissue was weighed out, placed into a 2 mL tube with 600 µL of RLT buffer with 1% β–mercaptoethanol and a 5 mm stainless steel bead (Qiagen, Hilden, Germany: #69989), then dissociated using a Qiagen TissueLyser II (Qiagen) with the following cycle: two min dissociation at 1800 oscillations/min, one min rest, two min dissociation at 1800 oscillations/min. Dissociated tissues were centrifuged at 13,000 rpm for 10 min at room temperature and cleared supernatant was transferred to a new 1.5 mL eppendorf tube. RNA extractions were performed using a Qiagen RNeasy Plus Mini Kit (Qiagen: #74134), according to the manufacturer's instructions (Qiagen: #79256). RNA was eluted in 30 µL of RNase/DNase free water and quantified by Nanodrop.

***fLX processing for viral titer quantification.*** Tissue pieces stored in RNALater were thawed at room temperature and then 22-40 mg of tissue were weighed out and placed into 600 uL of OptiMEM (ThermoFisher Scientific). Tissues were homogenized using a Qiagen TissueLyser II with two dissociation cycles, centrifuged and supernatant was transferred to a new 1.5 mL Eppendorf tube (see "*RNA extraction from fLX" section*). Tissue homogenates were then serial diluted ($10^{-1}$ – $10^{-6}$) and 300 uL of supernatant was plated on VeroE6 cells in 12-well plates ($2x10^5$

cells/well). Supernatant was incubated for 1 hour at 37°C to allow for viral adsorption then 1 mL of a 1:1 mixture of 2X DMEM 4% FBS 2% penicillin/streptomycin and 2.4% Avicel (Dupont) was overlaid onto each well. Plates were incubated for 72 hours at 37°C with 5% $CO_2$. Avicel was then removed, cells were fixed in 10% buffered formalin (ThermoFisher Scientific) for 1 hour, then stained with 0.1% crystal violet in 10% ethanol/$H_2O$ for 30 minutes before washing and quantification.

***Serum neutralization assay.*** One day prior to the experiment, 1 × 104 Vero E6 cells were plated into a 96-well plate. Serum was decomplemented at 56 °C for 30 min and an initial dilution of 1:10 was prepared in OptiMEM. Two-fold dilutions were subsequently prepared and mixed with rSARS-CoV-2 NL virus (MOI = 1) for 1 h at room temperature and then plated onto cells. After a 1-h incubation at 37 °C inoculum was removed and 200 µL of fresh DMEM containing 2% FBS and 1% penicillin/streptomycin was added. After a 24 h incubation at 37 °C with 5% CO2 media was removed and cells were fixed with 10% formalin for 1 h. A SARS-CoV-2 spike neutralizing antibody (Sino Biological Inc., Beijing, China; 2 µg/µL; Clone #57) was used as a positive control. Cells were washed with 1× PBS and 20 µM furimazine (MedChem Express, Monmouth Junction, NJ, USA) luciferin substrate was added onto cells. Cells were then imaged using an IVIS spectrum imager (PerkinElmer) and analyzed using LivingImage software (PerkinElmer). Titers were determined as the reciprocal of the highest dilution with >50% reduction of cytopathic effect.

***Quantification of peripheral human chimerism in BLT-L mice.*** 2–4x$10^6$ PBMCs of human or murine origin were isolated as described above and stained for 1 hour at 4°C in the dark with an antibody cocktail targeting human(h)CD45, mouse CD45, hCD3, hCD4, hCD8, hCD16, hCD19, hCD11c, hCD56 and hCD14. Following washing with FACS Buffer, cells were fixed with fixation buffer (1% (v/v) FBS, 4% (w/v) PFA in PBS) for 30 min at 4°C in the dark. Chimerism of all humanized mice was assessed by quantifying the following human populations: Human CD45+, human CD45+ murine CD45-; T-cells, CD45+ CD3+; CD4+ T cells, CD45+ CD3+ CD4+; CD8+ T cells, CD45+ CD3+ CD8+; CD45+ CD16+ leukocytes; B-cells, CD45+ CD19+; conventional dendritic cells, CD45+ CD11c+; NK/NKT cells, CD45+ CD56+; Monocytes, CD45+ CD14+.

***Antibody staining.*** After generation of single cell suspension, $5x10^5$ - $1x10^6$ cells were used for flow cytometry staining. Cells were centrifuged at 300 X *g* for 5 min at 4°C. The cell pellet was resuspended in a mix of 22.5 µL FACS buffer and 2.5 µL of FcX (Biolegend; #422302) and incubated for 10 min at room temperature. After blocking, 25 µL of antibody cocktail targeting hCD3, hCD20, hCD16, hHLA-DR, hCD45, hCD8, hCD4, hCD33, hCD45RA, hCD56, hCD14, mCD45, and containing a LIVE/DEAD viability dye (ThermoFisher Scientific) was added to each sample and incubated in the dark for 30 min at 4°C. After staining, 1 mL of FACS buffer was added to each sample, samples were centrifuged at 300 x g for 5 min, washed with 1 mL FACS buffer, centrifuged at 300 x g for 5 min, and then fixed in 200 µL 4% PFA for 30 min. After fixation cells were washed twice with 1 mL FACS buffer, resuspended in FACS buffer, and stored protected from light at 4°C until analysis. Human immune cell subsets were gated as follows: human CD45+, hCD45+ mCD45-; human CD3+, hCD45+ hCD3+; human CD4+, hCD45+ hCD3+ hCD4+; human CD8+, hCD45+ hCD3+ hCD8+.

***Flow cytometry analysis.*** For all flow cytometry experiments, flow cytometric analysis was performed using an LSRII Flow Cytometer (BD Biosciences) or Cytek Aurora instrument (Cytek Biosciences). Flow cytometry fluorophore compensation for antibodies was performed using an AbC™ Anti-Mouse Bead Kit (ThermoFisher Scientific). Flow cytometry data were analyzed using FlowJo software (TreeStar, Ashland, OR, USA).

***Histologic processing and analysis.*** Tissue samples were fixed for 72 h in 10% neutral buffered formalin, processed in a Tissue-Tek VIP-5 automated vacuum infiltration processor (Sakura

Finetek USA, Torrance, CA, USA), followed by paraffin embedding using a HistoCore Arcadia paraffin embedding station (Leica, Wetzlar, Germany). Generated formalin-fixed, paraffin-embedded (FFPE) blocks were sectioned to 5 µm using a RM2255 rotary microtome (Leica), transferred to positively charged slides, deparaffinized in xylene, and dehydrated in graded ethanol. Tissue sections were stained with hematoxylin and eosin for histologic examination, while unstained slides were used for immunohistochemistry. Qualitative and semi-quantitative histomorphological analyses were performed by a single board-certified veterinary pathologist (N.A.C.). An ordinal scoring system was developed to capture the heterogeneity of histologic findings in fLX. Individual histologic findings that were scored included: syncytial cells, hyaline membranes, intra-airspace neutrophils and necrosis, hemorrhage, edema, denuded pneumocytes, capillary fibrin thrombi, intermediate vessel fibrin thrombi and coagulative necrosis. The entire fLX was examined at 200x with a DM2500 light microscope (Leica) using the following criteria: 0 = not present, 1 = found in <5% of fields, 2 = found in >5% but <25% of fields, or 3 = found in >25% of fields. Several criteria were also restricted to 'not observed' (0) or 'observed' (1). Scores were combined to generate a cumulative lung injury score.

***Histological Antibodies.*** The following primary antibodies from Cell Signaling Technology (Danvers, MA, USA) were used: rabbit anti-human/mouse PAPRg (clone C26H12), rabbit anti-human CD4 (clone EP204), rabbit anti-human CD163 (clone D6U1J), rabbit anti-human CD3e (clone D7A6E), mouse anti-MHC class I (clone EMR8-5), and mouse anti-SARS-CoV-2 Spike (S) (clone E7U6O/2B3E5]). The secondary antibody used in this study included HRP Goat anti-Rabbit IgG (H&L) (Vector Laboratories, Burligame, CA, USA). For mouse derived primary antibodies, a linker antibody (Abcam) was used prior to application of the secondary antibody to prevent non-specific binding. DAB chromogen (Roche) and chromogens used for TSA-conjugated Opal 480, 520, 570, and 620 fluorophores (Akoya Biosciences, Marlborough, MA, USA) were utilized to develop immunohistochemical assays. The following anti-SARS-CoV-2 antibodies were used for immunohistochemistry: rabbit polyclonal anti-SARS-CoV Nucleoprotein (Novus Biological, Littleton, CO, USA), mouse monoclonal anti-SARS-CoV-2 Spike clone 2B3E5 (This antibody was used in this study as clone E7U60, which was the pre-production clone ID of clone 2B3E5; Cell Signaling Technology).

***Multispectral fluorescent imaging.*** Fluorescently labeled slides were imaged using either a Mantra 2.0TMor Vectra Polaris TM Quantitative Pathology Imaging System (Akoya Biosciences). To maximize signal-to-noise ratios, fluorescently acquired images were spectrally unmixed using a synthetic library specific for the Opal fluorophores used in each assay plus DAPI. An unstained fLX section was used to create an autofluorescence signature that was subsequently removed from multispectral images using InForm software version 2.4.8 (Akoya Biosciences).

***Transmission electron microscopy.*** Tissue samples were fixed for 72 h in a mixture of 2.5% glutaraldehyde and 2% formaldehyde in 0.1 M sodium cacodylate buffer (pH 7.4). Samples were then washed in 0.1 M cacodylate buffer and postfixed with 1% osmiumtetroxide (OsO4)/1.5% potassiumferrocyanide (KFeCN6) for 1 h at room temperature. After washes in water and 50 mM maleate buffer pH 5.15 (MB), the samples were incubated in 1% uranyl acetate in MB for 1 h, washed in MB and water, and dehydrated in grades of alcohol (10 min each: 50%, 70%, 90%, 2x10 min 100%). The tissue samples were then put in propyleneoxide for 1 h and infiltrated overnight in a 1:1 mixture of propyleneoxide and TAAB Epon. The following day the samples were embedded in fresh TAAB Epon and polymerized at 60°C for 48 h. Semi-thin (0.5 µm) and ultrathin sections (50-80 nm) were cut on a Reichert Ultracut-S microtome (Leica). Semi-thin sections were picked up on glass slides and stained with toluidine blue for examination by light microscopy to find affected areas in the tissue. Ultrathin sections from those areas were picked up onto formvar/carbon coated copper grids, stained with 0.2% lead citrate and examined in a JEOL

1200EX transmission electron microscope (JOEL, Akishima, Tokyo, Japan). Images were recorded with an AMT 2k CCD camera.

***Bulk RNA sequencing.*** Total RNA was processed from fLX as described above, and sent to BGI genomics (Hong Kong, China) for library preparation and sequencing (Pair-ends, 100 bp, 20M reads per sample). Raw FASTQ files were quality-checked with FastQC v0.11.7. Reads were found to be excellent quality and were aligned to the combined human (GRCh38, Ensembl 101) and mouse (GRCm38, Ensembl 101) genomes with STAR v2.7.1a (*2*) followed by quantification with featureCounts v1.6.2 (*3*). Quality was checked with MultiQC v1.6. All samples passed quality thresholds of >75% sequences aligned and >15 million aligned reads per sample. Significantly up- and downregulated genes were identified with DESeq2 v1.23.10 in R v3.6.0. (*4*). Each post-infection timepoint was compared to naïve: 2 dpi vs. naïve, and 12 dpi vs. naïve. P-values were FDR-adjusted, and log2 fold change was shrunk with the apeglm method. Significance was determined by an FDR-adjusted $p<0.01$ and a shrunken log2 fold change>2 or <-2. DESeq2 result were imported into Ingenuity Pathway Analysis (IPA; Service curated by Qiagen; Access provided through the Boston University Genome Science Institute) (*5*), and a canonical pathway enrichment analysis was performed using the default settings and the same differential expression thresholds as before (shrunken log2 fold change>2 or <-2 and FDR-adjusted p-value<0.01).

***Single-cell barcoding and sequencing.*** Following fLX processing into single-cell suspension as described above, cells were frozen down in a 90% FBS (Bio-Techne, R&D systems) 10% DMSO solution (ThermoFisher Scientific) and kept at -80°C. Four to five days following freezing, cells were thawed, and viability was assessed using Trypan blue (Fisher Scientific). Samples with ≥90% viability were then processed using the Chromium Next GEM Single Cell 3' GEM, Library & Gel Bead Kit v3.1, as per manufacturer instructions, and single-cell barcoding was performed using a Chromium instrument (10x genomics) located in the NEIDL BSL-3. Reverse transcription of RNAs was performed in the BSL-3 using a thermocycler (Applied Biosciences), and cDNA was then removed from containment. Full-length, barcoded cDNAs were then amplified by PCR to generate sufficient mass for library construction. Enzyme fragmentation, A tailing, adaptor ligation and PCR were then performed at the Boston University single-cell sequencing core to obtain final libraries containing P5 and P7 primers used in Illumina bridge amplification. Size distribution and molarity of resulting cDNA libraries were assessed via Bioanalyzer High Sensitivity DNA Assay (Agilent Technologies, USA). All cDNA libraries were sequenced on an Illumina NextSeq 500 instrument at the Boston University microarray and sequencing core according to Illumina and 10x Genomics guidelines with 1.4-1.8pM input and 1% PhiX control library spike-in (Illumina, USA).

***Single-cell RNA sequencing analysis.***

Sequencing data was processed via 10X Genomics Cell Ranger 6.0.2. A custom reference was built from the combined human (GRCh38, Ensembl 101) and mouse (GRCm38, Ensembl 101) genomes. FASTQ files were generated from raw Illumina BCL files using *cellranger mkfastq*. FASTQ files were aligned to the custom reference and quantified with *cellranger count*. Quantification outputs were loaded and analyzed in R with the Seurat package (4.0.2, (*6*)). Cells with nFeature_RNA between 200 and 7500, and MT.percent<= 50 were analyzed. The dataset was normalized and scaled, and cell cycle scores (S+G2M) and nCount_RNA were regressed. Seurat's RunPCA function was used to determine the number of PCA used for subsequent analysis. FindClusters function was used to cluster cells. RunUMAP function was used for UMAP visualization. A human score was calculated using all human genes, to discriminate between mouse and human cells (**Fig. S3A-C**). Human cells were kept for subsequent analysis. The following ("dimension";"resolution") parameters were used to cluster the whole human dataset (15;0.1), epithelial cells (10;0.3), mesenchymal (10;0.4), endothelial (10;0.5), T cell/ILC (12;0.4), and Myeloid cells (10;0.2). B cell, endothelial cell, epithelial cell, mesenchymal cell, and mast cell clusters were identified by scoring using gene lists defined for lung subsets in the COVID-19 atlas of ToppCell (**Fig. S3D**) (*7*). For chondrocytes, which are not present in this dataset, we used the following gene list *ACAN, ITGA10, CYTL1, HAPLN1, SNORC, COL2A1, COL9A2, COL11A2, SOX9, SOX6, ZBTB16, FOXC1, and GATA2*. For T cells and ILC, a manually curated gene list based on ToppCell was used. Consensus signature genes from LGEA LungMAP (*8*) were used to validate each cluster. Differentially expressed genes between clusters and conditions were identified using FindMarkers. Only genes showing expression in at least 10% of cells in one of the groups compared, p_val_adj < 0.05, and abs(fold change)>2 were considered.

***Sequencing of SARS-CoV-2 virus from fLX.*** Viral sequences were amplified using a modified version of the ARTIC3 tiled amplicon sequencing protocol (*9, 10*) and sequenced on an Illumina MiSeq V3 600 cycle kit. Viral RNA was extracted from fLX as described above in "*RNA extraction from fLX*". cDNA was synthesized from the viral RNA using a mix of random hexamer and oligo-dT primers (IDT) and SSIV reverse transcriptase (Thermo Fisher). Tiled amplicons were amplified using Q5 Hot Start polymerase (NEB) and the pooled primers described in ARTIC3 (*9, 10*). Illumina sequencing adaptors were ligated onto the amplicons using the NEBNext Adaptor kit for Illumina (NEB). The amplicons were then pooled, cleaned up, and size selected using AMPureXP beads (Beckman Coulter). The pooled library was quantified using Tape Station (Agilent) and Qbit (Thermo Fisher) kits. The library was sequenced on a MiSeq with a V3 600 cycle kit (Illumina). Sequencing reads were filtered, trimmed, and mapped to the D-WA reference sequence (Geneious). Variants were called using the Geneious variant caller, with a minimum frequency threshold of 25%. Each positive variant was visually inspected.

### SUPPLEMENTAL FIGURES

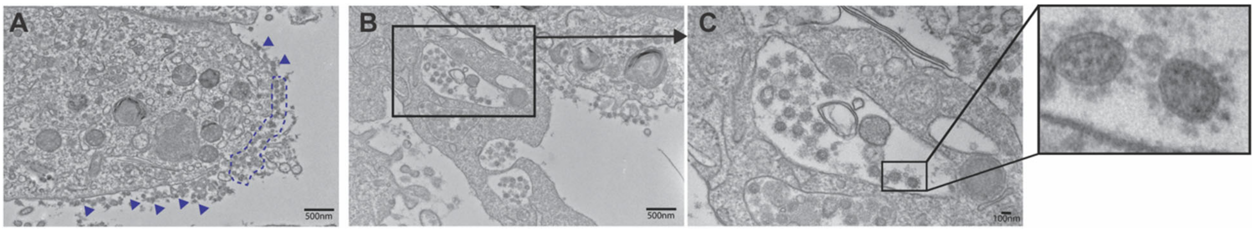


**Figure S1. BLT-L mice are susceptible to SARS-CoV-2 infection. (A-C)** Transmission electron microscopy (TEM) of fLX tissue sections extracted from BLT-L mice at 2 dpi illustrating virus particles at the cell surface as indicated by the blue arrows and blue dotted line **(A)** and viral particles in AT2 cells as evident by presence of lamellar bodies **(B,C**). Scale bars indicated on image.

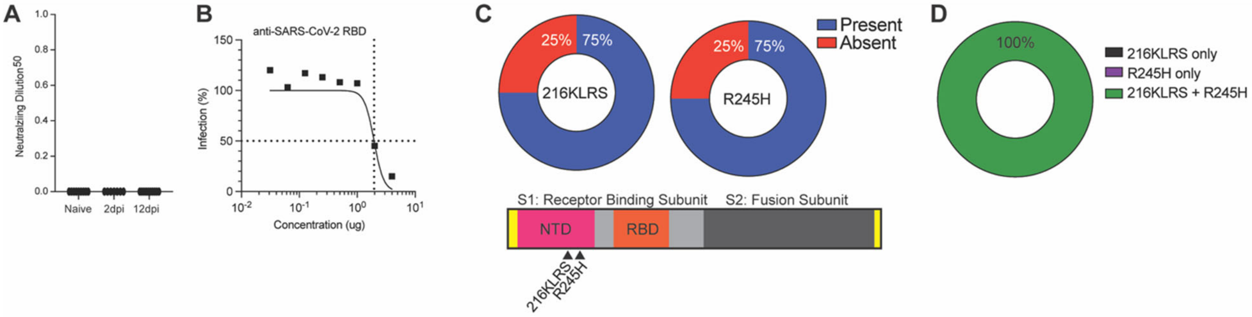


**Figure S2. SARS-CoV-2 undergoes viral adaptation within fLX. (A)** Neutralizing efficacy (ND50 values) of serum extracted from naïve or infected mice BLT-L mice (2 and 12 dpi). Assay was performed using VeroE6 cells and a recombinant WA-1 SARS-CoV-2 virus expressing NanoLuc. **(B)** Titration of the neutralizing activity of an anti-RBD antibody (serving as positive control) against WA-1 SARS-CoV-2 virus expressing NanoLuc. **(C-D)** SARS-CoV-2 virus isolated from 2 dpi fLX was deep sequenced and assessed for mutations relative to the inoculation (WA-1) strain. **C Upper panel**: Pie chart depicting the percentage of fLX with virus containing 216KLRS insertion and R245H non-synonymous mutation. **C Lower Panel**: Schematic representation of

SARS-CoV-2 spike protein highlighting the location of mutations. *Not to scale.* **D:** Pie chart depicting the percentage of fLX with virus that showed signs of co-evolution of 216KLRS and R245H.

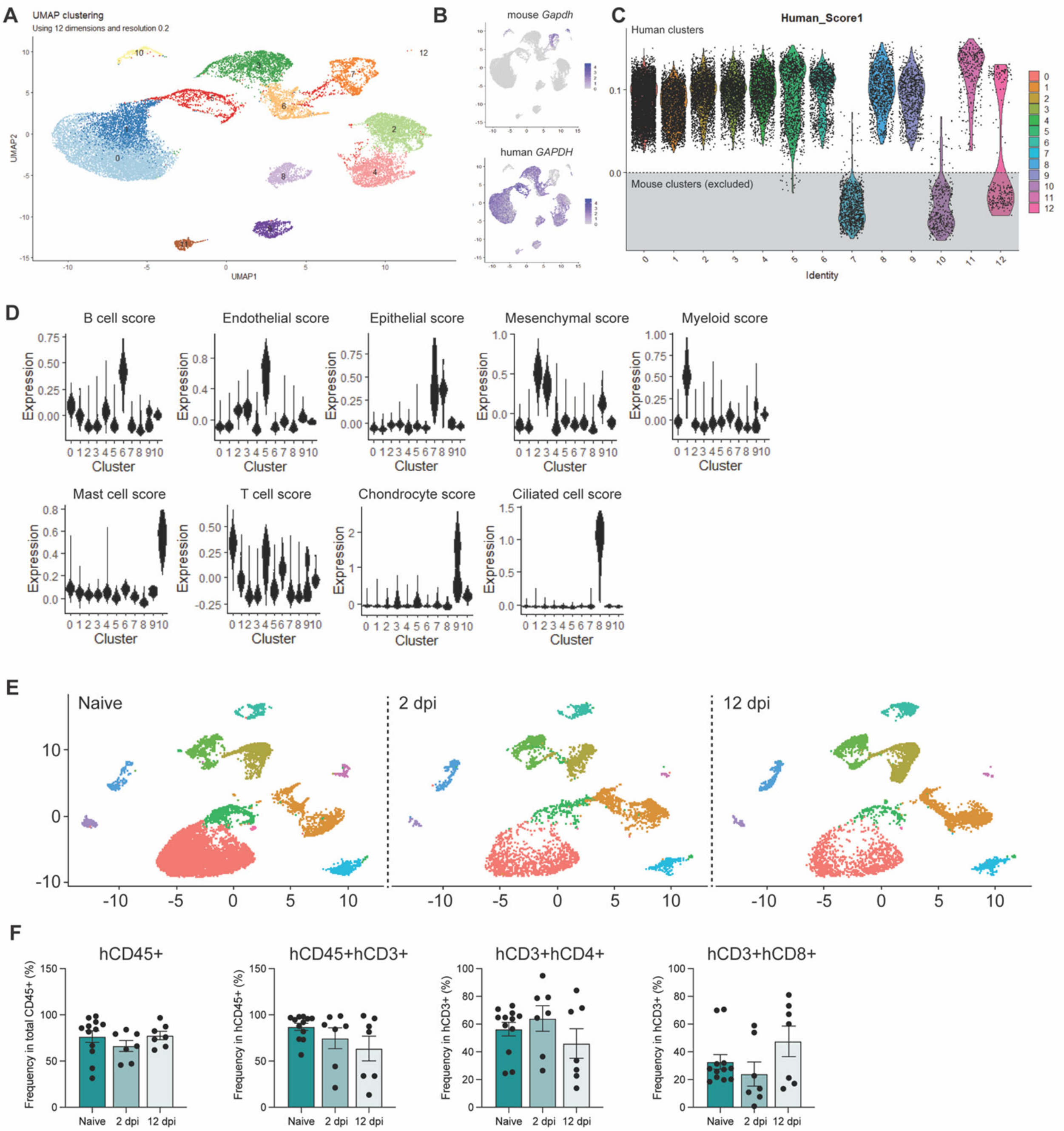

**Figure S3. Single-cell RNA sequencing analysis of fLX upon SARS-CoV-2 infection. (A-E)** Single cell RNA sequencing was performed on naïve fLX and infected fLX (at 2- and 12 dpi). Sequencing reads were aligned to a combined human, mouse and SARS-CoV-2 viral genome. **(A)** UMAP plot clustering on all cell (human and mouse) populations detected. **(B)** Expression of mouse (top) and human *GAPDH* (bottom) in all clusters. **(C)** A human score was applied to each cluster to identify human clusters (above dotted line) and mouse clusters (below dotted line; gray zone). Mouse clusters were removed from downstream analysis. **(D)** Cell scoring system applied to human cell clusters to classify each cluster. **(E)** UMAP clustering of human cells separated by time point (naïve: left, 2 dpi: center, and 12 dpi: right). **(F)** Flow cytometric analysis showing frequency of hCD45+, hCD3+, hCD4+, and hCD8+ cells among PBMCs extracted from the blood of naïve and infected BLT-L mice at 2-, 6-, and 12 dpi.

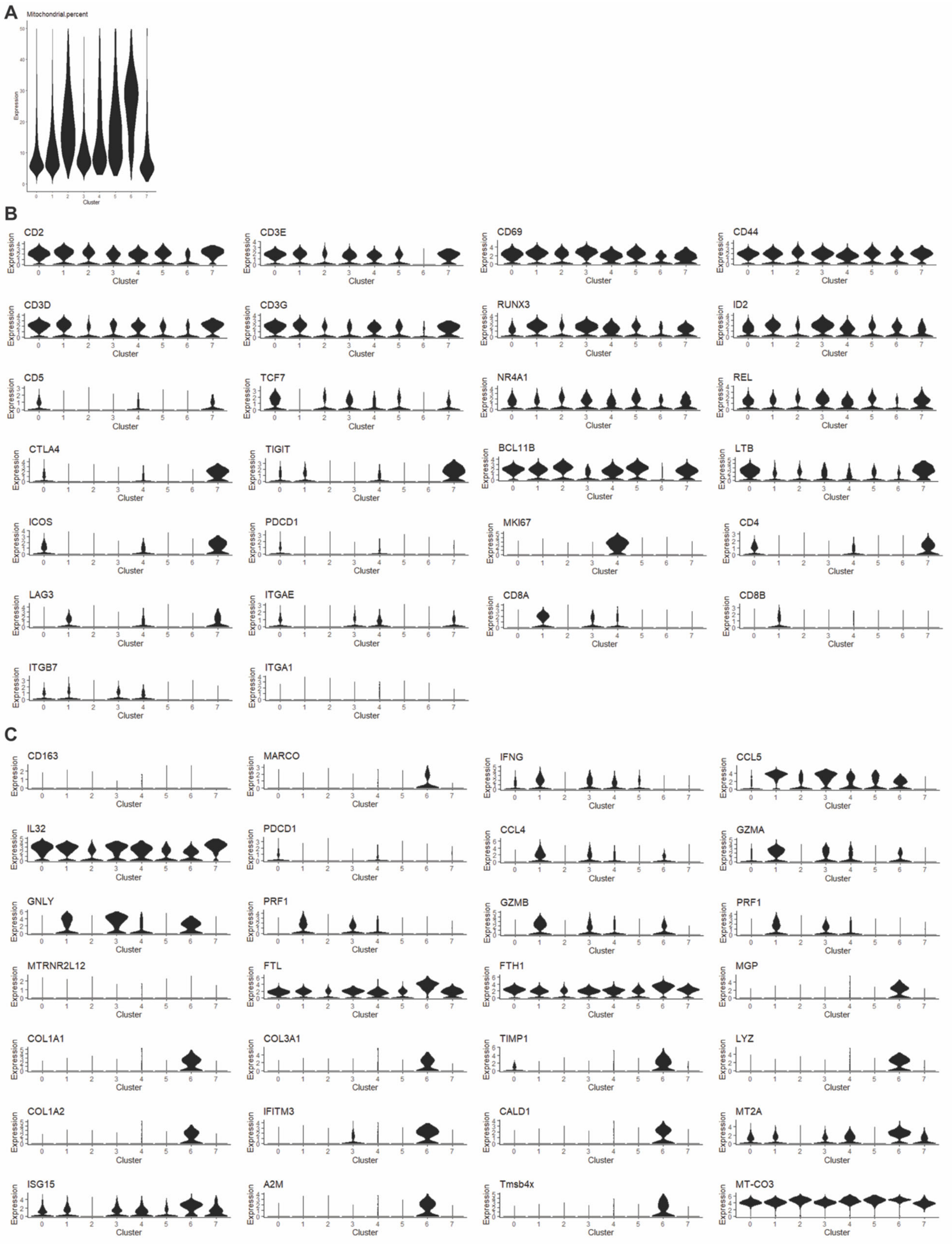
A
Mitochondrial.percent
Expression
Cluster
B
CD2
CD3E
CD69
CD44
CD3D
CD3G
RUNX3
ID2
CD5
TCF7
NR4A1
REL
CTLA4
TIGIT
BCL11B
LTB
ICOS
PDCD1
MKI67
CD4
LAG3
ITGAE
CD8A
CD8B
ITGB7
ITGA1
C
CD163
MARCO
IFNG
CCL5
IL32
PDCD1
CCL4
GZMA
GNLY
PRF1
GZMB
PRF1
MTRNR2L12
FTL
FTH1
MGP
COL1A1
COL3A1
TIMP1
LYZ
COL1A2
IFITM3
CALD1
MT2A
ISG15
A2M
Tmsb4x
MT-CO3

**Figure S4. Gene signature of macrophage-like T cells. (A)** Mitochondrial gene expression among the different T cell sub-clusters. **(B)** Expression of T-cell associated genes among the different T-cell sub-clusters. **(C)** Expression of myeloid/macrophage-associated genes among the different T-cell sub-clusters.

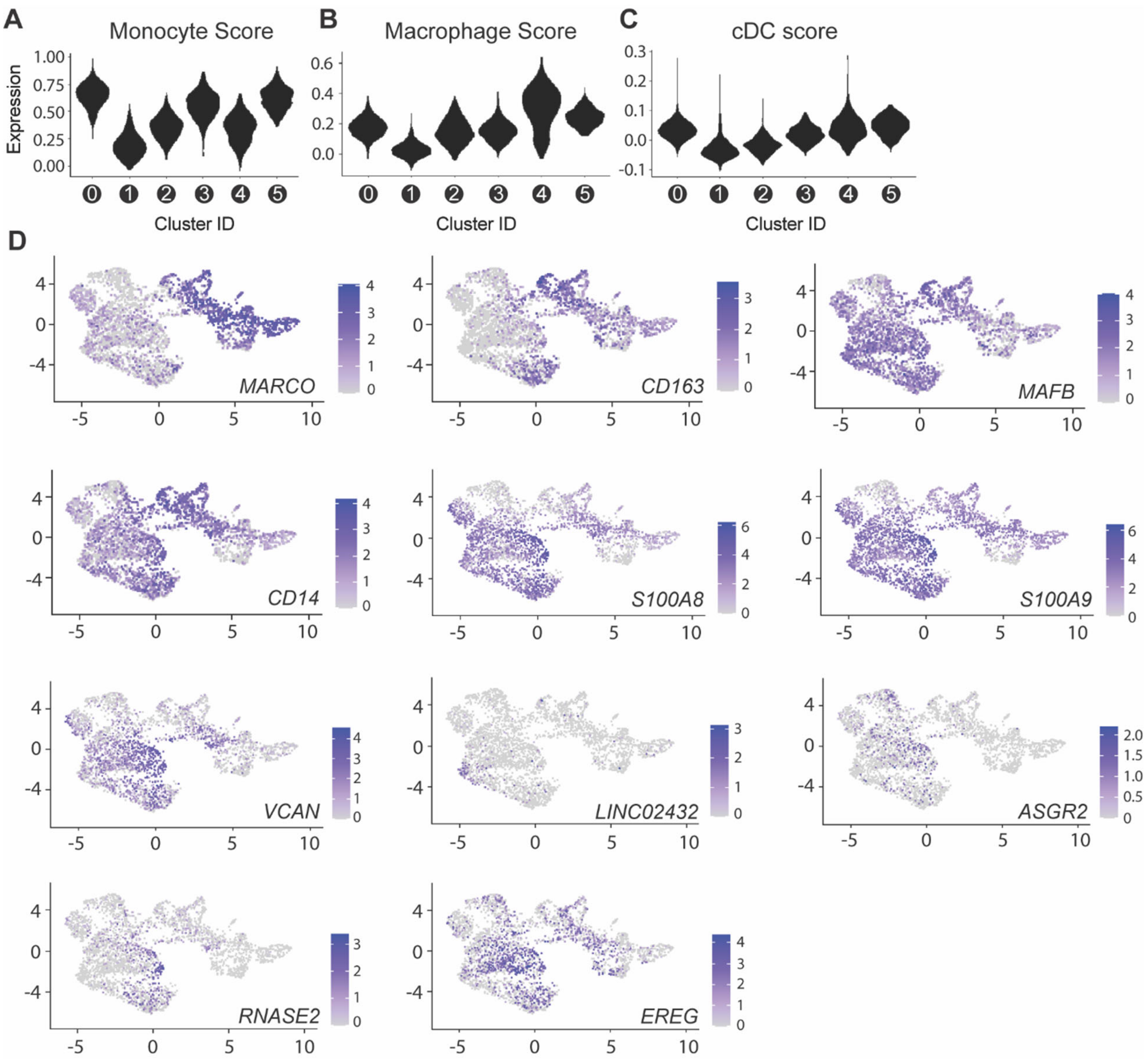


**Figure S5. Gene signature of myeloid and endothelial sub-clusters. (A-C)** Violin plot displaying monocyte **(A)**, macrophage **(B)** and dendritic cell **(C)** gene signature score for each myeloid sub-cluster. **(D)** UMAP plots representing the expression of several myeloid markers that were employed to further define the identity of the different myeloid sub-clusters.

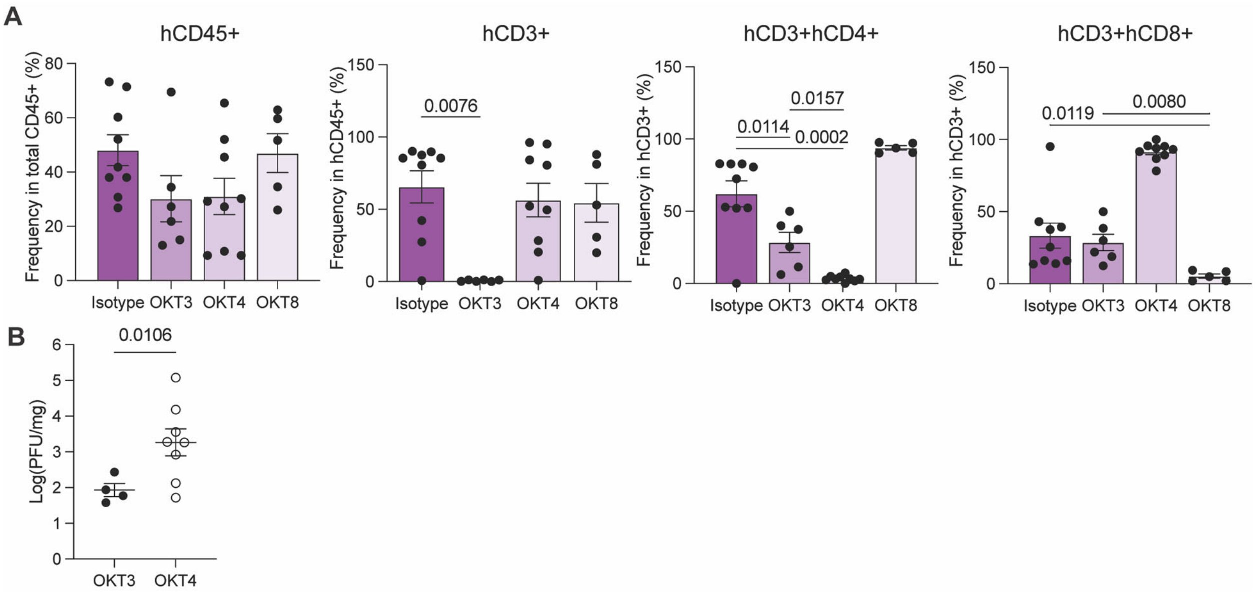


**Figure S6. Assessment of systemic depletion efficiency and analysis of viral titers in mice with abrogated viral clearance mechanisms. (A)** Flow cytometric analysis showing the frequency of hCD45+, hCD3+, hCD3+ hCD4+, and hCD3+ hCD8+ cells among PBMCs extracted from the blood of BLT-L mice treated with an isotype antibody, OKT3, OKT4 or OKT8 antibody. Error bars represent mean ± SEM. *One-way ANOVA. P-value indicated on graph.* **(B)** Viral titer (log(PFU/mg)) of CD3+ cell (OKT3) and CD4+ cell-depleted (OKT4) fLX that were positive for viral infection. *Unpaired, non-parametric T-test. P-value indicated on graph.*

### SUPPLEMENTAL TABLES

| Sample | Mutations | Notes |
|---|---|---|
| **2 dpi - 1** | **Spike 216KLRS insertion 99%** | There is a deletion at spike 221, but there is very low coverage (only 2x) |
| | **Spike R245H - 100%** | |
| **2 dpi - 2** | orf1ab K1051 insertion - 28% | no mutations in spike, but spike had no coverage at AA sites 216 and 245 |
| **2 dpi - 3** | **Spike 216KLRS insertion 100%** | |
| | **Spike R245H - 100%** | |
| **2 dpi - 4** | **Spike 216KLRS insertion 91%** | |
| | **Spike R245H - 100%** | |
| **2 dpi - 5** | **Spike 216KLRS insertion 81%** | |
| | **Spike R245H - 100%** | |
| **2 dpi - 6** | **Spike 216KLRS insertion 96%** | |
| | **Spike R245H - 99%** | |
| | NS6 S43P - 30% | |
| **2 dpi - 7** | **Spike 216KLRS insertion 98%** | |
| | **Spike R245H - 100%** | |
| **2 dpi - 8** | orf1ab V1866 del (frameshift) 35% | low coverage at AA sites 216 and 245, but there is 1 read supporting the presence of both mutations |
| | orf1ab V3048L 33%, | |
| | NS3 - F168 del (frameshift)- 47%, | |
| **WA1 positive stock** | Spike E96A - 51% | |
| | Spike 678-690 deletion 87% | |
| | NS7b 43 - NS8 1 deletion 50% | |

**Table S1.** SARS-CoV-2 mutations identified among viral reads isolated from fLX at 2 dpi and from our concentrated viral stock (i.e., WA1 positive stock used to inoculate fLX), as compared to the Wuhan-1 consensus sequence.
